# Genetic transformation of *Gardnerella* species and characterization of vaginolysin and sialidase mutants

**DOI:** 10.1101/2025.05.09.653190

**Authors:** Amy K. Klimowicz, Erin M. Garcia, Kimberly K. Jefferson, Joseph P. Dillard

## Abstract

Bacterial vaginosis (BV) is the most prevalent vaginal disorder in women of childbearing age and causes pregnancy complications including preterm birth. Species of *Gardnerella* increase just prior to the onset of symptoms and are considered to play major roles in the development and transmission of BV. However, *Gardnerella* species have remained genetically intractable, limiting investigations of their virulence mechanisms. Here we describe methods for genetic manipulation of *Gardnerella*. Through trial and error we optimized methods for electrotransformation of *Gardnerella* and created methods for making mutations and complements. We mutated the gene for the toxin vaginolysin (*vly*) in *G. vaginalis* and the gene for sialidase *nanH3* in *G. pickettii*. A *vly* point mutant was tested in human cervix tissue and found to lack lytic activity. The *nanH3* mutant lost sialidase and mucus degradation activity. Overall, this genetic toolkit opens a door for molecular characterization of *Gardnerella* and its mechanisms in BV.

**Teaser:** Genetic studies are now possible in a causative agent of bacterial vaginosis, and mutants were made lacking the toxin or mucus-degrading enzyme.

## Introduction

Genetic manipulation is a fundamental process in the study of bacterial pathogenesis. The abilities to make mutations and complements are necessary for definitive identification of virulence genes, and the ability to make unmarked mutations is preferable in these studies (*1*). Certain bacterial pathogens were considered genetically intractable for many years, such as *Mycobacterium tuberculosis* or *Chlamydia trachomatis*, and studies of their virulence mechanisms necessarily focused on epidemiology, immunology, cell biology in the host, and proteomic or transcriptomic analyses (*2, 3*). Those classic cases were eventually resolved by the discovery of effective transduction, natural transformation, and chemical transformation methods, greatly facilitating the identification of virulence factors (*4-7*). *Gardnerella* species have similarly been considered genetically intractable, and like *M. tuberculosis* and *C. trachomatis*, *Gardnerella* have an unusual cell envelope. Similar to other Actinobacteria, *Gardnerella* are generally considered Gram-positive, but stain Gram-variable because of their unusual and incompletely characterized envelope (*8*).

The *Gardnerella* genus was originally thought to be made up of a single species, *Gardnerella vaginalis*. Many studies of *Gardnerella* diversity divided the isolates by clade, ecotype, or *cpn60* type, and recent genomic studies have determined that there are at least thirteen species (*9-12*). Six species have been formally named: *G. vaginalis, G. piotii, G. swidsinskii, G. leopoldii, G. greenwoodii*, and *G. pickettii* (*12, 13*). The differences in presence or absence of suspected virulence genes between species have led some researchers to propose that some *Gardnerella* species are pathogens while others are commensals or that different *Gardnerella* species play different roles in causing the dysbiosis of bacterial vaginosis (BV) (*14-16*). Other studies suggest that *Gardnerella* species are pathogenic when present with other *Gardnerella* species, i.e., that a mix of two or more *Gardnerella* species is needed to initiate the dysbiosis of BV (*17*). The genetic methods described here will allow researchers to identify virulence factors in the various *Gardnerella* species and dissect the roles of these factors in development of the dysbiosis and the associated pathology.

BV affects nearly one-third of women of reproductive age, causing problems for sexual and reproductive health (*18-20*). BV in pregnant women greatly increases the risk of preterm birth, and preterm birth is the leading cause of infant mortality (*21*). Furthermore, women with BV are more likely to acquire a range of sexually-transmitted infections, including gonorrhea, chlamydia, and HIV, and are more likely to transmit HIV (*22, 23*). BV is difficult to cure. Although *Gardnerella* species are sensitive to antibiotics, only 60-70% of patients are cured following primary treatment, and a significant portion of patients who respond to treatment will relapse within a period of months (*24*). Thus, better methods are needed to combat this common and poorly-controlled disease. Identification of crucial virulence factors would allow for the development of vaccines or targeted therapeutics. As with other dysbioses, microbiota transplant may represent a viable treatment (*25*). While BV patients usually have very high levels of *Gardnerella* bacteria present, nearly all women harbor *Gardnerella* as vaginal colonizers (*15, 26*). Thus, the identification of harmless *Gardnerella* species using molecular studies or the creation of harmless *Gardnerella* species by deletion of virulence genes would allow for the creation of vaginal microbiota consortia that could be used for microbiota transplant or for prevention of infection with pathogenic *Gardnerella*.

Several factors are predicted to act in pathogenesis of *Gardnerella* species (*27*). *Gardnerella* are proposed to cause the dysbiosis of BV by adhering to epithelial cells, building a biofilm that multiple BV-associated bacteria grow on, lysing human cells, degrading host molecules, cross-feeding other BV-associated bacteria, degrading IgA, and reducing the inflammatory response (*24, 28-33*). Putative adhesins and biofilm factors include Tad pili, sortase-dependent pili, and proteins predicted to bind extracellular matrix and serum molecules including fibronectin, albumin, and collagen (*34-36*). Most *Gardnerella* produce the toxin vaginolysin which lyses red blood cells, epithelial cells, and neutrophils (*37*). Such lysis would release numerous host molecules that the bacteria could use for nutrition and for supplying nutrients to other BV-associated bacteria. Glycogen degradation and protein degradation are described as supplying *Gardnerella* and the other bacteria in the biofilm with nutrients. Sialidase is present in some *Gardnerella* species and is proposed to act in the degradation of mucins. *Gardnerella* sialidase has been shown to act on IgA, potentially leading to IgA destruction (*38*). Sialidase activity correlates with BV and is used as a diagnostic factor (*39*). Other glycosylases such as fucosidase and beta-galactosidase are likely also involved in mucus degradation. It has not previously been possible to test the roles of these factors in infection models. In this study, we made mutations in the vaginolysin gene and a sialidase gene and demonstrated effects on cell lysis in human cervix tissue and degradation of human cervical mucus.

## RESULTS

### Preparation of electrocompetent cells and transformation of *G. vaginalis* with oligonucleotides

To determine if DNA could be transformed into *G. vaginalis* by electroporation, we attempted oligo-mediated recombination (*40, 41*). *G. vaginalis* ATCC 14018 was selected for these experiments, and the large colony variant was used due to its robust growth (*35*). Cells were grown in liquid culture to early log-phase (OD_600_ ∼0.35), chilled on ice, harvested by centrifugation, and washed with 10% glycerol. The final cell density in this suspension was approximately 2.0x10^9^ CFU/ml. For oligo-mediated recombination, we designed a mutagenic 44-base oligonucleotide that would introduce a dinucleotide substitution (AA to CG) in the *rpsL* gene, conferring streptomycin resistance. In other bacteria, transformation efficiency can be increased by using levels of oligonucleotides that saturate the host exonucleases (*41*). Thus, we used a relatively high concentration of oligo; *G. vaginalis* cells were mixed with 10 µg of oligo in ice-cold cuvettes and electroporated at 2.2 kV. Cells were allowed to recover overnight in supplemented brain heart infusion medium (BHI-YDS) before plating on agar medium containing streptomycin. After two days, Str^R^ colonies were obtained at frequencies of approximately 10^-6^ Str^R^ CFU/total CFU (**Fig. 1A**). Sequencing of the *rpsL* gene in Str^R^ isolates showed that the 2-base mutation had been incorporated.

**Figure 1.**
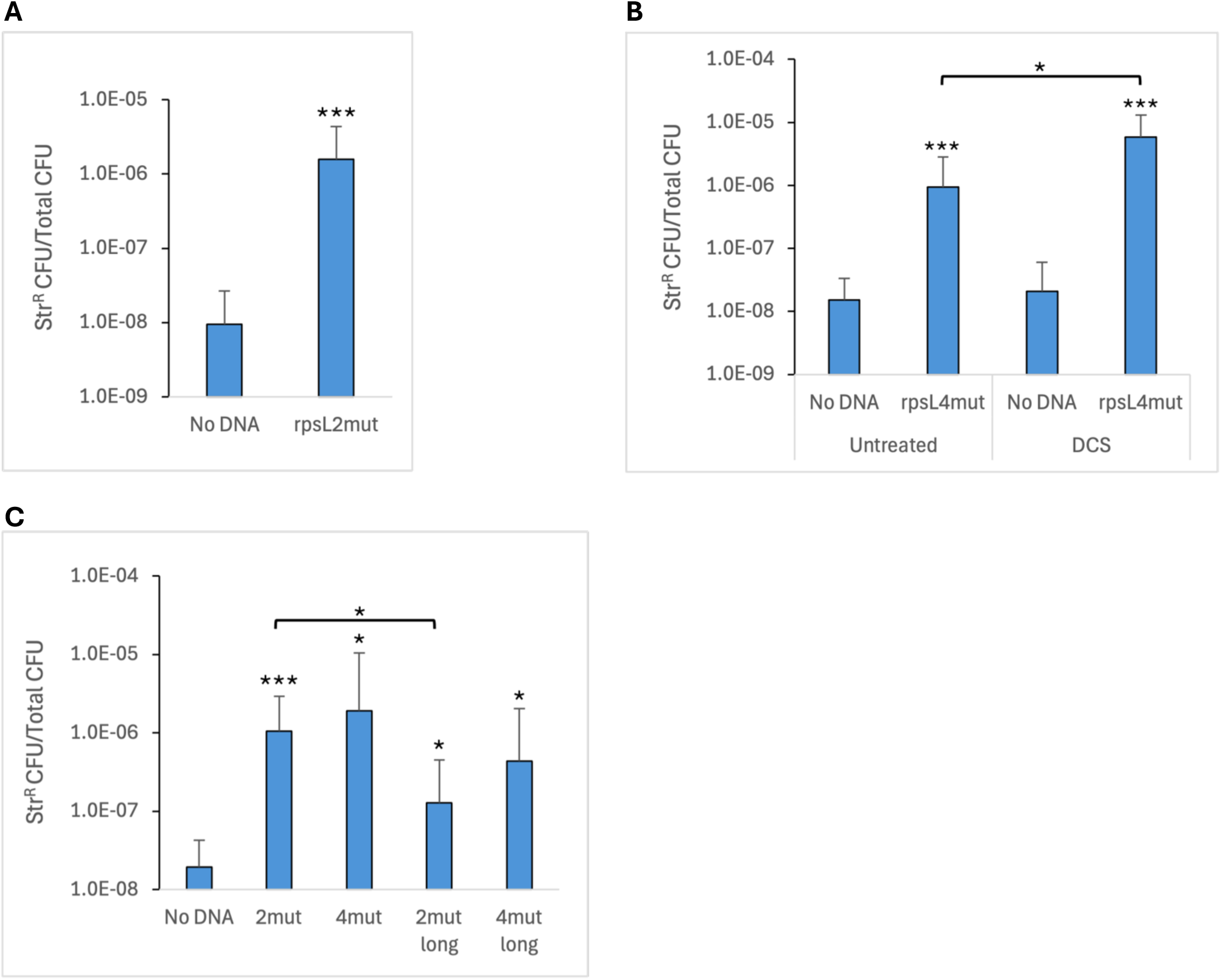
Oligo-mediated recombination in *G. vaginalis* ATCC 14018. (A) Competent cells were prepared without DCS treatment and transformed with 10 µg of oligo rpsL2mut. The No DNA control shows the frequency of spontaneous Str^R^ observed in cells electroporated without DNA. (B) Competent cells were prepared with or without DCS treatment and transformed with 10 µg of oligo rpsL4mut. (C) Competent cells were prepared with DCS treatment and transformed with diHerent *rpsL* mutagenic oligos. The oligos were 44- or 84-nt in length and contained either a 2- or 4-nt change (2mut or 4mut) from the wild-type sequence. The geometric means and standard deviations of three independent assays is shown. Statistical significance in (A) and (B) was assessed by the Student’s *t*-test. Significance in (C) was assessed by either the Student’s *t*-test, or the Welch’s *t*-test for 4mut versus No DNA and 4mut long versus No DNA. Significance is shown relative to the respective No DNA control or between oligos as indicated by brackets. * *P* < 0.05, ** *P* < 0.01, *** *P* < 0.001.

Once oligo-mediated recombination had been achieved, we altered a variety of parameters to determine the impact on transformation efficiency. Harvesting cells at a later point in growth (OD_600_ ∼0.5) resulted in a 4-fold reduction in the number of transformants compared to cells from early log-phase. Reducing the voltage of electroporation from 2.2 kV to 1.8 kV resulted in half as many recombinants. Increasing the amount of oligo from 10 µg to 50 µg did not impact the number of transformants, suggesting that 10 µg was a saturating amount of oligonucleotide. No Str^R^ colonies were obtained if the electroporation was done with a room temperature cuvette. Also, no transformants were recovered if the cells were allowed to recover for only 4 hours. Using frozen and thawed cells, rather than freshly prepared cells, reduced efficiency by one third.

### Growth of *G. vaginalis* with D-cycloserine increases transformation efficiency

We investigated whether growing the cells in the presence of potential cell wall-weakening compounds could enhance the transformation frequency. We grew ATCC 14018 in the presence of glycine, Tween 80, lysozyme, penicillin, DTT, or lithium acetate, and we electroporated them with an *rpsL* oligo containing 4 base changes (TAAG to ACGC), conferring streptomycin resistance. None of these treatments increased the efficiency of transformation with the *rpsL* oligo. We also tried growing cells in the presence of sucrose and washing with a sucrose-based buffer, but again we did not observe an increase in transformation frequency. However, addition of a sub-lethal concentration of D-cycloserine (DCS) to the culture had a positive effect on transformation efficiency. ATCC 14018 was grown to an OD_600_ = 0.2-0.25, at which time DCS was added to a concentration of 250 µg/ml, and the culture was grown for another 1.5 hours. Following growth with DCS, we observed that cultures exhibited a reduced OD_600_ compared to untreated cultures. Electroporation of DCS-treated cells with the *rpsL* oligo resulted in a 6-fold increase in Str^R^ frequency compared to non-treated cells (**Fig. 1B**). Therefore, we included this treatment in subsequent competent cell preparations. DCS is an antibiotic that inhibits alanine racemase and D-Ala D-Ala ligase, both of which are involved in peptidoglycan synthesis (*42*). It is likely that our treatment of *G. vaginalis* with DCS reduces the extent of crosslinking between peptidoglycan strands and thereby weakens the barrier function of the cell envelope.

### Longer oligos do not result in higher transformation frequencies

In *E. coli*, the DNA mismatch repair system can correct mismatches which naturally occur during replication, but it also represents a barrier to the introduction of point mutations. However, mismatch repair can be overcome by introducing consecutive mutations or C-C mismatches (*43*). We compared transformation frequency with the 44-base *rpsL* oligo containing mismatches in 4 consecutive nucleotides (TAAG to ACGC) to the original oligo that has 2 nucleotide changes (AA to CG). The 44-base oligo with the 4-base mutation resulted in a 1.8-fold higher average transformation frequency compared to the oligo with the 2-base mutation, but this apparent difference was not statistically significant (**Fig. 1C**). We also tested two longer oligos, both 84 nucleotides, that contained the same 2-base or 4-base mutations, to determine whether a longer homologous region would increase transformation efficiency. However, electroporation with the longer oligo resulted in an 8-fold lower transformation frequency compared to the corresponding shorter oligo for the 2-base mutation oligos in three independent assays (**Fig. 1C**). A similar trend was seen for the 4-base mutation oligos, but the difference was smaller and did not reach the level of significance.

### Creation of a *vly* (vaginolysin) mutant using circular, methylated DNA

After successful transformation with the *rpsL* oligos, we attempted to transform various larger pieces of DNA into *G. vaginalis*. We tried chromosomal DNA from *Gardnerella* strains that naturally carry a *tetM* marker. We tried linear constructs which we built carrying an *ermC* or *tetM* marker inserted into the *vly* (vaginolysin) gene which we had cloned from strain ATCC 14018. None of these forms of DNA worked for transformation. Next, we considered that small plasmids might work for transformation. No plasmids have been isolated from *Gardnerella* spp. (*9*). Therefore, we tried to obtain an insertion-duplication mutant by integration of an *E. coli* plasmid into the *G. vaginalis* chromosome, selecting for erythromycin resistance encoded on the plasmid. We cloned an approximately 0.5 kb fragment internal to the vaginolysin gene into *E. coli* plasmid pIDN1, and electroporated 2 μg of resulting plasmid pAKK134 into ATCC 14018 (**Fig. 2A**). Our first attempts to obtain plasmid transformants were unsuccessful. Erm^R^ isolates were obtained at a frequency no different from a no-DNA control, and PCR analysis determined that these isolates did not contain the plasmid construct. However, the report of a HaeIII-like type II restriction-modification (RM) system in ATCC 14018 that cleaves at unmethylated GGCC sequences suggested that this endonuclease activity might be preventing plasmid transformation (*44*). There are seven HaeIII sites in pAKK134. To avoid restriction of plasmid DNA, we treated pAKK134 with HaeIII methyltransferase and repeated the electroporation. We obtained Erm^R^ colonies at a frequency of 1.6x10^-7^, over 50-fold higher than the frequency of spontaneous Erm^R^ colonies obtained in the negative control (2.8x10^-9^). PCR analysis with one primer that anneals to the chromosome and one that anneals to the plasmid confirmed that nine Erm^R^ isolates from the plasmid transformation all contained pAKK134 integrated at *vly* (**Fig. 2B**). A hemolysis assay with concentrated cell supernatants from the wild type or one of the insertion-duplication mutants showed that the mutant was reduced in hemolytic activity compared to the wildtype (**Fig. 2C**). The small colony variant (Sm) of wild-type strain ATCC 14018 produces large amounts of vaginolysin, while the large colony variant (Lg) produces much less (**Fig. 2C**, (*35*)). The *vly* mutant strain AKK107 exhibited only background levels of hemolytic activity against human red blood cells whether the Sm or Lg variant was used (**Fig. 2C**).

**Figure 2.**
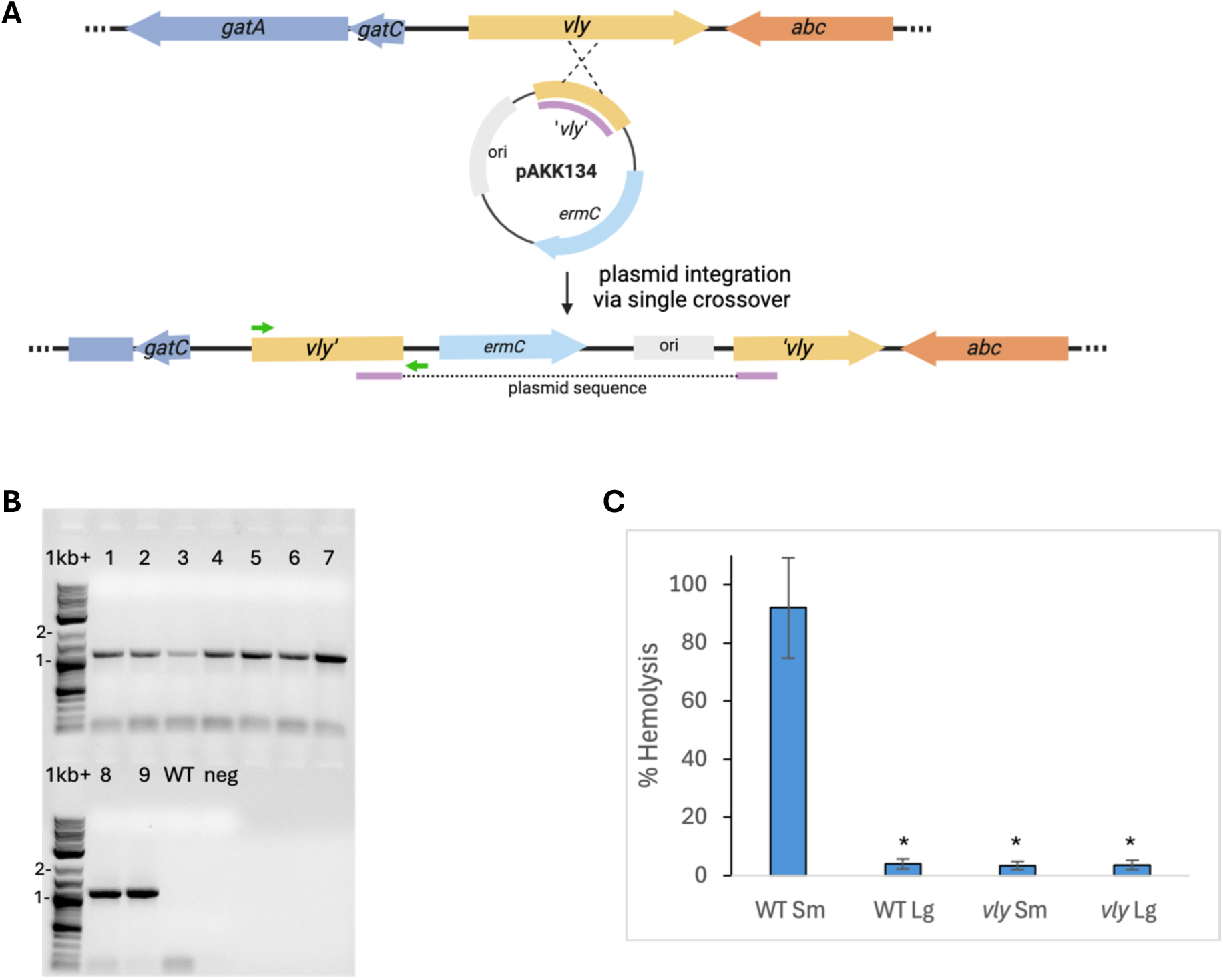
Creation of a *vly* insertion-duplication mutation eliminates hemolysis activity in *G. vaginalis*. (A) Schematic of the *vly* region of ATCC 14018 and pAKK134 integrated via a crossover in *vly*. (B) PCR analysis of nine Erm^R^ isolates from the 14018 x pAKK134 transformation with primers (green arrows) to the chromosome (vly-F) and to the vector (pmob-R), producing a 1.1-kb product if pAKK134 is integrated at *vly*. 1kb+ = NEB 1kb plus DNA ladder, WT = ATCC 14018, neg = No DNA control. (C) Hemolysis assay of small (Sm) and large (Lg) colony variants of ATCC 14018 and the insertion-duplication mutant, AKK107. Supernatants from overnight growth were concentrated with 30-kDa cut-oH columns, and dilutions were incubated with a 1% suspension of human red blood cells. After removal of unlysed cells, the OD_415_ of the supernatants was measured. The average percent hemolysis is shown for three biological replicates. *, significantly diHerent from WT Sm at *P* < 0.05 by the Welch’s *t*-test.

### A *haeIIIR* mutant is transformable with unmethylated plasmid

In subsequent transformations with untreated pAKK134 and methylated pAKK134, we found that sometimes transformants were obtained with untreated plasmid, but the frequency with methylated plasmid was on average 37-fold higher than with untreated plasmid (**Fig. 3A**). We next constructed an insertion-duplication mutant to knock out the HaeIII-like endonuclease gene in ATCC 14018 to examine whether transformation of the mutant with an untreated plasmid could be obtained at a frequency similar to that with a methylated plasmid. ATCC 14018 was transformed with Me-pAKK139 (‘*haeIIIR*’), resulting in Erm^R^ insertion-duplication mutant AKK110. We then transformed AKK110 with methylated or untreated plasmid pAKK149, which is identical to pAKK134 but with the *ermC* replaced by *aad9*, encoding spectinomycin adenyltransferase. Transformation of AKK110 with untreated and HaeIII-methylated pAKK149 resulted in Spc^R^ colonies at a relatively high frequency for this procedure, approximately 1x10^-6^ for both conditions (**Fig. 3B**). The similar frequencies for both methylated and untreated plasmids indicates that the HaeIII-like restriction-modification system was inhibiting plasmid transformation in wild-type ATCC 14018.

**Figure 3.**
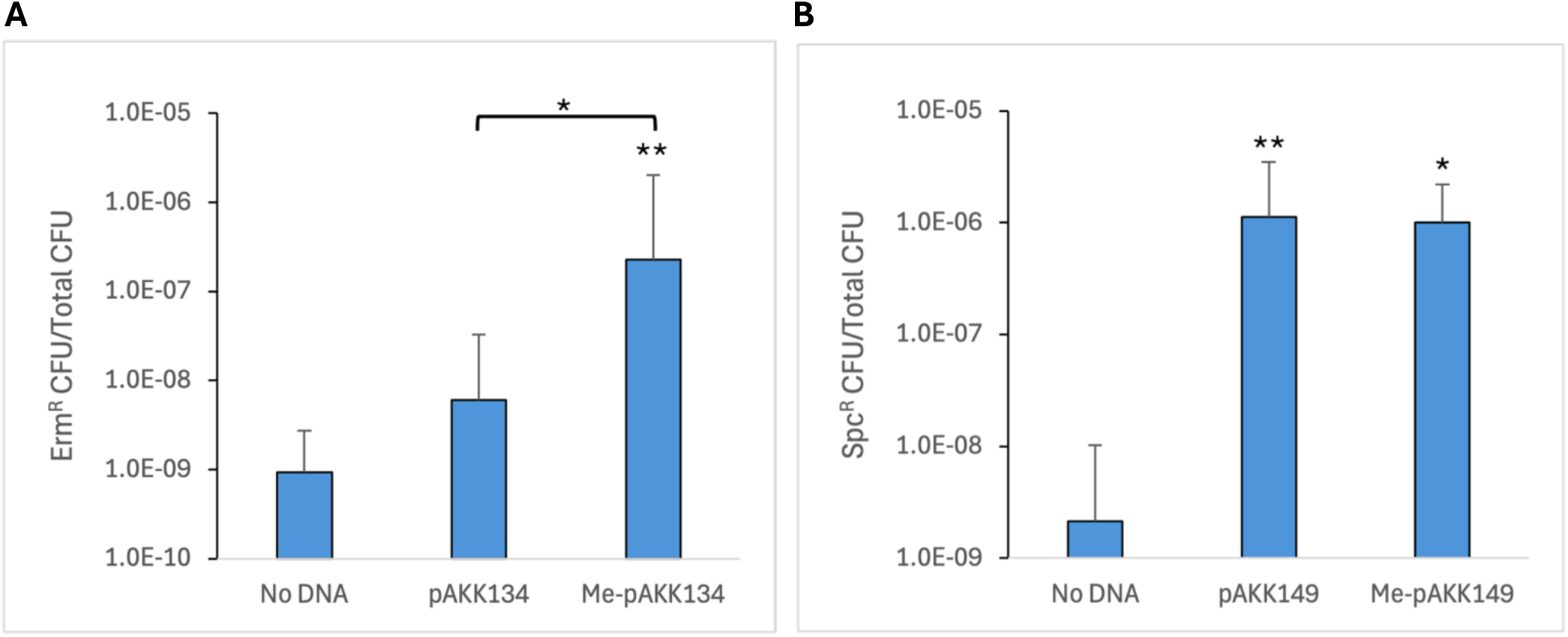
The HaeIII-like restriction-modification system in *G. vaginalis* ATCC 14018 presents a barrier to plasmid transformation. (A) Competent cells of ATCC 14018 were transformed with 2 µg of untreated pAKK134 (‘*vly*’) or 2µg of pAKK134 that had been treated with HaeIII methyltransferase. The No DNA control shows the frequency of spontaneous Erm^R^ mutants obtained. The geometric mean and standard deviation of four independent assays is shown. Statistical significance was assessed by the Student’s *t*-test. (B) Transformation of insertion-duplication mutant AKK110 (*haeIIIR*) with 2 µg of untreated or methylated pAKK149 (‘*vly*’). The No DNA control shows the frequency of spontaneous Spc^R^ mutants obtained. The geometric mean and standard deviation of three independent assays is shown. Statistical significance was assessed by the Student’s *t*-test, or the Welch’s *t*-test for Me-pAKK149 versus No DNA. Significance is shown relative to the No DNA control or between plasmids as indicated by brackets. * *P* < 0.05, ** *P* < 0.01.

### Circular DNA constructs transform better than linear constructs

To determine if our initial attempts at transformation might have failed due to using linear DNA, we compared transformation frequencies for constructs that were either circular or linearized plasmids. Plasmid pAKK111 contains 1.9kb of *G. vaginalis* chromosomal DNA with a *tetM* marker inserted near the middle of the cloned region. The *tetM* gene is inserted between 984 bp of *vly* region DNA and 912bp of the downstream gene, which encodes an ABC transporter. Linear pAKK111 was produced by digestion with restriction enzyme SphI, which cuts in the plasmid backbone, outside the region homologous to the chromosome. Tetracycline resistant transformants were obtained at a 7-fold higher frequency with the circular plasmid as compared to the linearized plasmid (**Fig. 4**). These data suggest that linear DNA may be more readily degraded than circular DNA prior to recombination or that single-crossover recombination occurs at higher frequency than double-crossover recombination.

**Figure 4.**
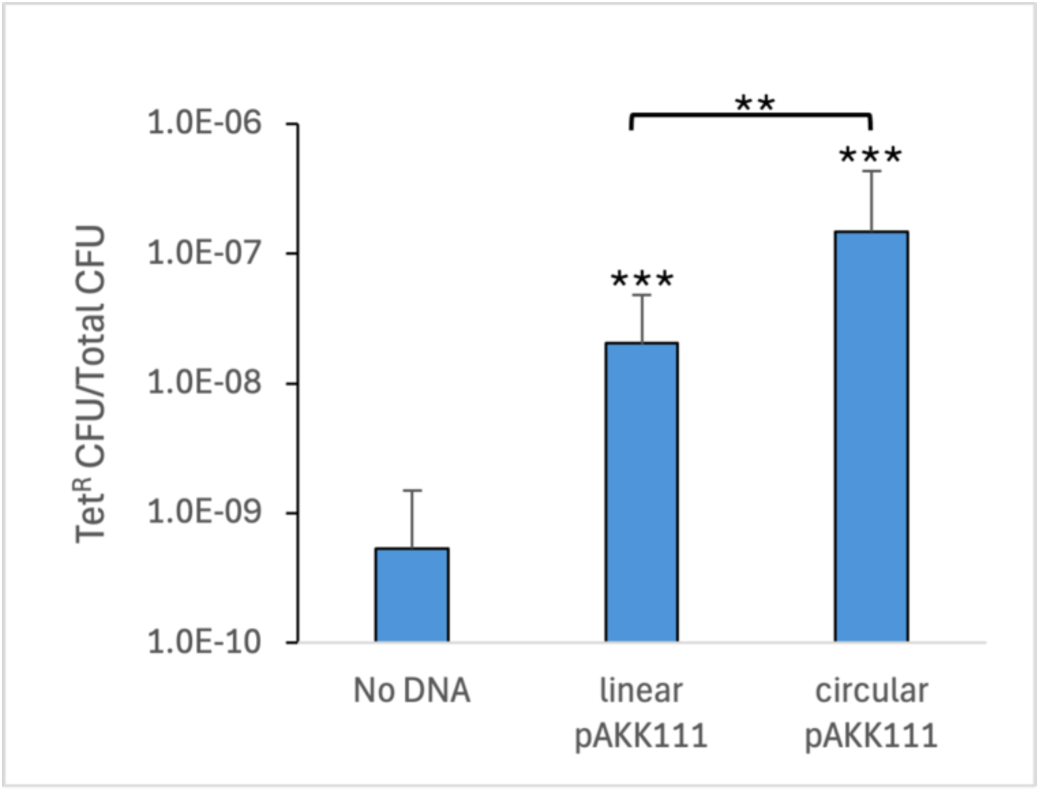
Circular constructs transform *G. vaginalis* better than linear constructs. Transformation of insertion-duplication mutant AKK110 (*haeIIIR*) with ∼2 µg of linearized or circular pAKK111 (Tet^R^). The No DNA control represents the limit of detection, as no colonies were obtained. The geometric mean and standard deviation of three independent assays is shown. Statistical significance was assessed by the Student’s *t*-test. Significance is shown relative to the No DNA control or between plasmids as indicated by brackets. ** *P* < 0.01, *** *P* < 0.001.

### Frequency of plasmid integration and excision correlates with length of homologous DNA

To examine how the frequency of plasmid integration might vary with the length of homologous DNA, we designed a series of constructs that contained an internal fragment of *G. vaginalis lacZ* that was 0.2-, 0.55-, 1-, or 1.8-kb in length. Plasmid integrants that interrupted the *lacZ* gene were selected for using erythromycin resistance encoded on the plasmid. We found that transformation frequency correlated with the size of the insert, ranging from an average of 5.0x10^-9^ with the 0.2-kb insert to 2.8x10^-7^ with a 1.8-kb insert (**Fig. 5**). Plasmid integrant colonies appeared white on media containing X-Gal, whereas the wild-type colonies were blue. To determine how stable the insertions were, we measured how frequently blue colonies arose after overnight growth of the mutants in non-selective liquid media. Restoration of the wild-type *lacZ* gene would occur in isolates that underwent a second recombination that excised the plasmid. The frequency of blue colonies varied inversely with the size of the duplication, ranging from an average of 2.3x10^-5^ blue colonies/total colonies with the 0.2-kb insert to 4.8x10^-3^ blue colonies/total colonies with the 1.8-kb insert. This is likely an overestimation of the excision frequency since excision occurring in colonies during early growth on the plate would also generate blue colonies. Nevertheless, given the relatively high frequency of plasmid excision in insertion-duplication mutants, we sought a method that would allow us to construct unmarked gene deletions and point mutations.

**Figure 5.**
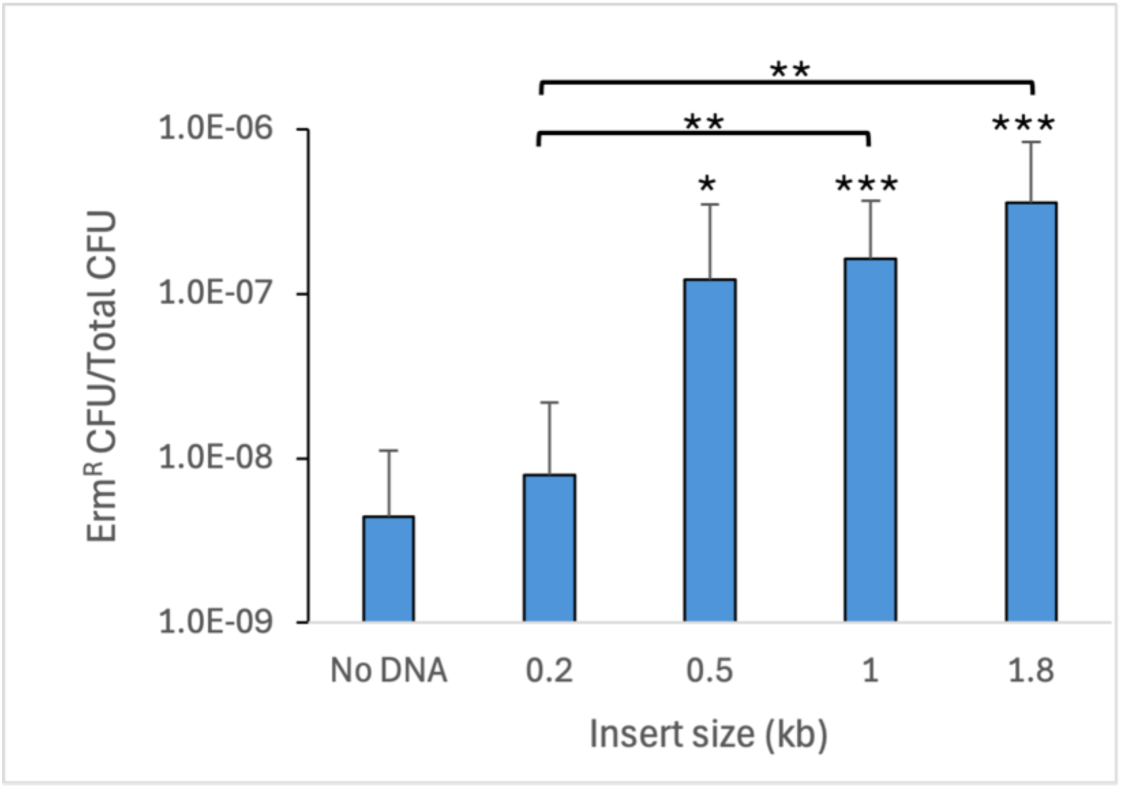
Length of homologous DNA correlates with the frequency of plasmid integration. ATCC 14018 was transformed with a series of plasmids that contain *ermC* for selection and an internal fragment of *lacZ*, ranging in size from 0.2-1.8 kb, for homologous recombination with the chromosome. The No DNA control shows the frequency of spontaneous Erm^R^ isolates obtained. The geometric mean and standard deviation is shown for three (0.2 kb, 0.5 kb) or four (No DNA, 1 kb, 1.8 kb) independent assays. Statistical significance was assessed by the Student’s *t*-test. Significance is shown relative to the No DNA control or between plasmids as indicated by brackets. * *P* < 0.05, **, *P* < 0.01, ***, *P* < 0.001.

### Construction of a positive/negative selection plasmid

In order to introduce unmarked mutations into *Gardnerella,* we developed an insertion plasmid that contains the *ermC* gene for selection of plasmid integration (positive selection), and a mutated *pheS* gene for the selection of plasmid loss (negative selection). Use of this positive-negative selection vector allows one to change specific bases in a gene or delete a specific gene using a two-step procedure. We employed *pheS* containing a double substitution as a counter-selectable marker. The *pheS* gene encodes the α subunit of phenylalanine-tRNA synthetase. In *E. coli,* PheS with an A294G mutation misincorporates the toxic phenylalanine analog 4-chloro-DL-phenylalanine (4CP) into proteins, leading to cell death (*45*). An improved *pheS* allele containing two mutations, T251S A294G, was engineered which allowed selection on rich media and with lower concentrations of 4CP (*46*). Alignment of the PheS proteins from *E. coli* and ATCC 14018 revealed that ATCC 14018 contained the conserved Thr and Ala residues present in *E. coli*. Therefore, we constructed a plasmid containing ATCC 14018 *pheS* with the double substitution (*pheS*_mut2_), preceded by the *G. vaginalis* promoter for the gene encoding glyceraldehyde-3-phosphate dehydrogenase (P_gap_), which would ensure a high level of expression of *pheS*_mut2_. We cloned this *pheS*_mut2_ into plasmid pIDN1, resulting in positive/negative selection plasmid pAKK163.

### Construction and characterization of a *vly* point mutation strain

To test whether pAKK163 could be used to create an unmarked mutation, we first constructed a point mutation in *vly* in ATCC 14018. We cloned a 0.7-kb fragment of the vaginolysin gene that spanned amino acids 39-271 and contained a nonsense mutation at E155 (GAA to TAA), resulting in plasmid pAKK165 (**Fig. 6A**). Following electroporation of the methylated plasmid into ATCC 14018, we obtained Erm^R^ transformants that had integrated the plasmid at *vly*, and they exhibited diminished hemolysis on HBT agar. We grew the integrant on medium without Erm to allow for a second recombination event and loss of the plasmid. We plated dilutions of the culture on medium with and without 1 mM 4CP, and we obtained 4CP^R^ colonies at a frequency of 4.7x10^-4^. Ninety-three percent of the colonies were also sensitive to Erm, demonstrating that the *pheS*_mut2_ provided efficient counter selection against the plasmid. Colonies from the 4CP plate were patched to HBT agar to screen for recombinants that had maintained the nonsense mutation and appeared nonhemolytic after overnight growth. Approximately 50% of the patches appeared reduced in hemolysis compared to the wild type. Sequencing of the *vly* gene in a nonhemolytic isolate (AKK124) confirmed the presence of the nonsense mutation (**Fig. 6B**).

**Figure 6.**
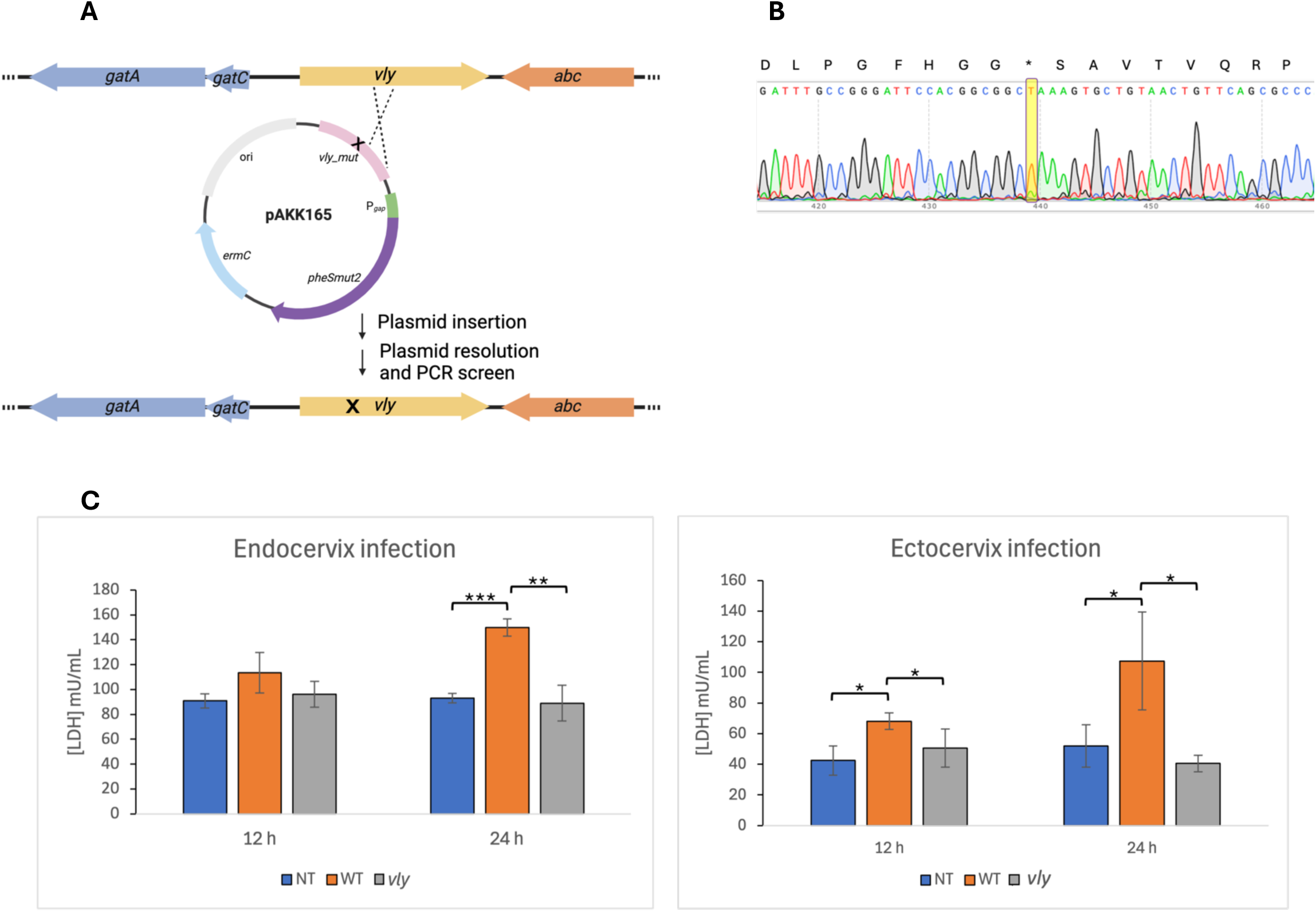
Construction and characterization of a *vly* point mutant of ATCC 14018 using a positive/negative selection plasmid. (A) Schematic depicting construction of a *vly* point mutant. Plasmid pAKK165 contains *ermC* for positive selection (plasmid integration) and *pheS_mut2_* for counter selection (plasmid excision). The 0.7-kb fragment of *vly* contains a Glu155* mutation, indicated by the “X.” (B) DNA sequencing confirmed that the single nucleotide change that results in a premature stop (GAA to TAA; E155*) in vaginolysin was present in mutant AKK124. The nt change is highlighted in yellow. (C) LDH levels released from endocervical (left panel) and ectocervical (right panel) tissues that were inoculated with media alone (NT, no treatment), ATCC 14018 (WT) or AKK124 (*vly*). LDH was quantified after 12 and 24 hr. The mean ± standard deviation of three (endocervix) or four (ectocervix) independent assays is shown. Statistical significance for endocervix infection was assessed by the Student’s *t*-test. For ectocervix infection significance was assessed by the Student’s *t*-test, or the Welch’s t-test for 24 hr WT versus the *vly* mutant. * *P* < 0.05, ** *P* < 0.01, ***, *P* < 0.001.

To quantify the cytotoxicity of the wild type and *vly* point mutant on cervical cells, we infected human ectocervical and endocervical tissue explants with the bacteria and measured the level of lactate dehydrogenase, a cytosolic enzyme that is released when cells are damaged. The LDH level from the wild type-infected tissues increased over time and exhibited statistically higher LDH levels than the mutant after 12 and 24 hr of infection of the ectocervical tissue (**Fig. 6C**, right panel), and after 24 hr infection of the endocervical tissue (**Fig. 6C**, left panel). The LDH activity measured for the *vly* mutant was no different from untreated tissue at 12 or 24 hr (**Fig. 6C**).

### Construction of a *vly* deletion strain

While working with derivatives of the positive/negative selection plasmid pAKK163, we found that the plasmid sometimes recombined at *pheS* in the *G. vaginalis* chromosome instead of at our target gene. Therefore, to prevent recombination from occurring at *pheS*, we recoded the *pheS_mut2_* gene, substituting *G. vaginalis* codons with those optimized for *Bifidobacterium longum*, a related bacterial species. This process resulted in the positive/negative selection construct pAKK174. We used this new vector to delete the entire vaginolysin coding region. We designed a plasmid that fused approximately 0.5 kb of DNA upstream and downstream of *vly* cloned in pAKK174, resulting in pAKK176 (**Fig. 7A**). Upon transformation of ATCC 14018, we obtained Erm^R^ transformants that had pAKK176 integrated in the region near *vly* at a frequency of 1.4x10^-8^, approximately 1 log higher than the frequency of spontaneous Erm^R^ colonies. After growing one of these isolates without Erm, the culture was plated on medium containing 4CP. Isolates that grew on 4CP and failed to grow on Erm-containing medium were patched to HBT to screen for those that had retained the *vly* deletion, evidenced by diminished hemolysis (**Fig. 7B**). PCR analysis of one of these non-hemolytic mutants confirmed it lacked *vly* (**Fig. 7C** and **7D**).

**Figure 7.**
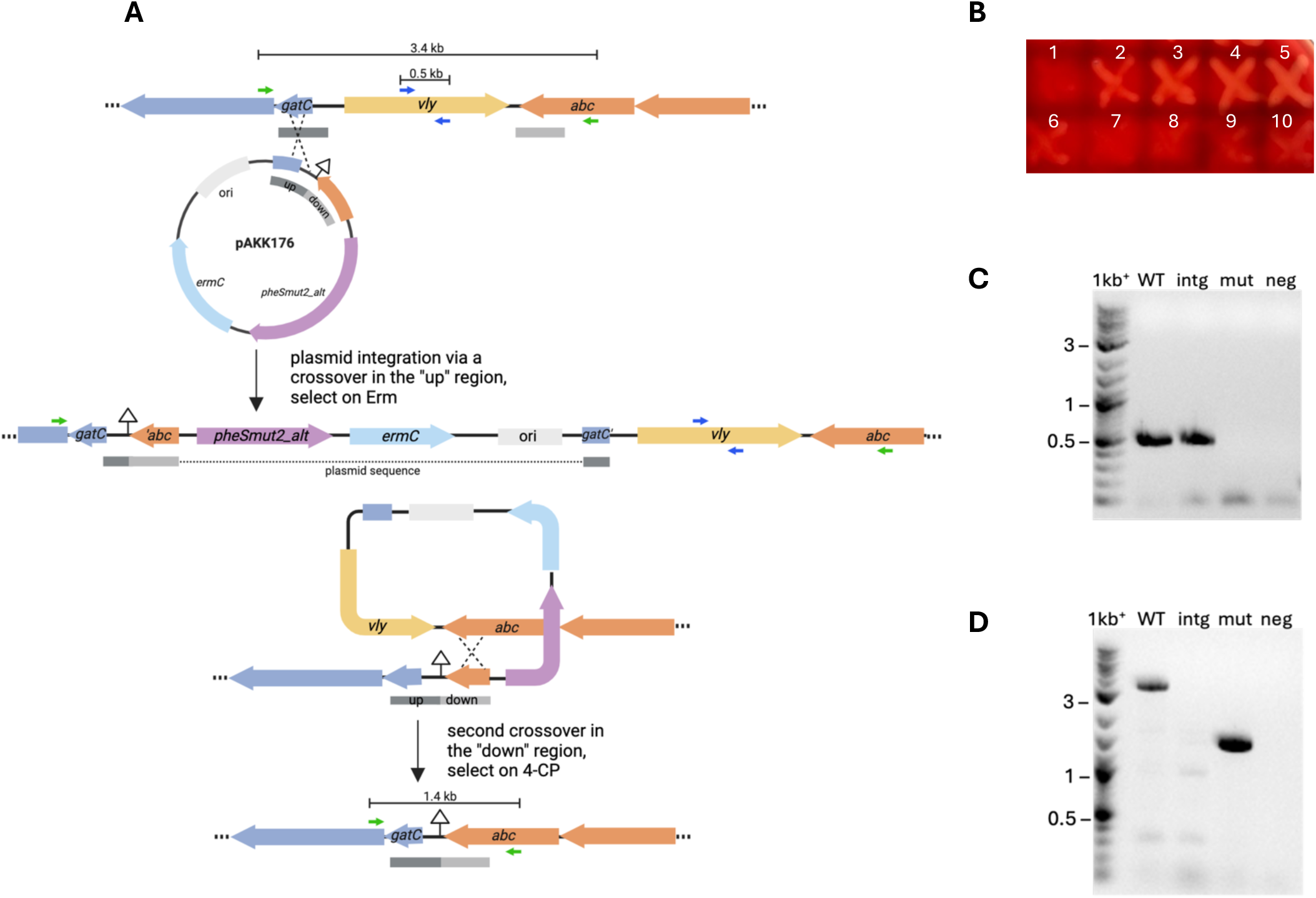
Markerless deletion of *vly* in *G. vaginalis* ATCC 14018 using a positive/negative selection strategy. (A) Schematic depicting the *vly* region in ATCC 14018 and plasmid pAKK176 used in the two-step gene replacement. pAKK176 contains fragments flanking *vly* designated “up” and “down.” In the first step, transformants that have integrated the plasmid are selected for by plating on media with Erm. In this example, integration occurred via a single crossover in the “up” region of the chromosome. In the second step, growth of the integrant without Erm, followed by plating on media containing 4-chlorophenylalanine (4CP), selected for isolates that had undergone a second recombination event and lost the plasmid. In this example, the second crossover occurred in the “down” region, resulting in deletion of *vly*. (B) Isolates that were resistant to 4CP and sensitive to Erm were patched to HBT agar to screen for recombinants that exhibited reduced hemolysis. Patches 2-5 appeared hemolytic, while patches 1 and 6-10 appeared to have diminished activity. The patch labeled “1” was analyzed by PCR. (C) and (D) PCR analyses confirming the *vly* deletion. (C) PCR with primers that are internal to *vly* (blue arrows in A; vly_mid-F/vly_screen-R). (D) PCR with primers that anneal outside of the “up” and “down” flanking regions (green arrows in A; vly_up_screen-F/Gv_abc-R). Lanes: 1kb+ = DNA ladder; WT = ATCC 14018; intg = pAKK176 integrant; mut = *vly* deletion mutant AKK132; neg = No DNA control. Note that the PCR extension time in D was not suHicient to allow amplification of the 7.6-kb product from the integrant.

### Complementation of the *vly* deletion mutant

To identify a site in the *G. vaginalis* genome that could be used to express genes for complementation, we scanned for a region that had a pair of convergent genes and a conserved chromosomal organization across different *Gardnerella* species. The region containing *pknB*, encoding a serine/threonine protein kinase, and *srtE*, coding for a class E housekeeping sortase, met these criteria. We designed a construct that contained approximately 0.8 kb of the 3’ end of *pknB*, and 0.8 kb of the 3’ end of *srtE,* along with a multiple cloning site (MCS) between these two gene fragments. To drive constitutive expression of the gene of interest, we included the promoter region from *rpsB* of ATCC 14018. Restriction sites on either side of the *rpsB* promoter (NcoI/AgeI) allow for the possibility of cloning in a different promoter. The complementation construct, pAKK186, has a spectinomycin resistance gene on the vector and the *ermC* gene within the region that will be incorporated into the chromosome (**Fig. 8**).

**Figure 8.**
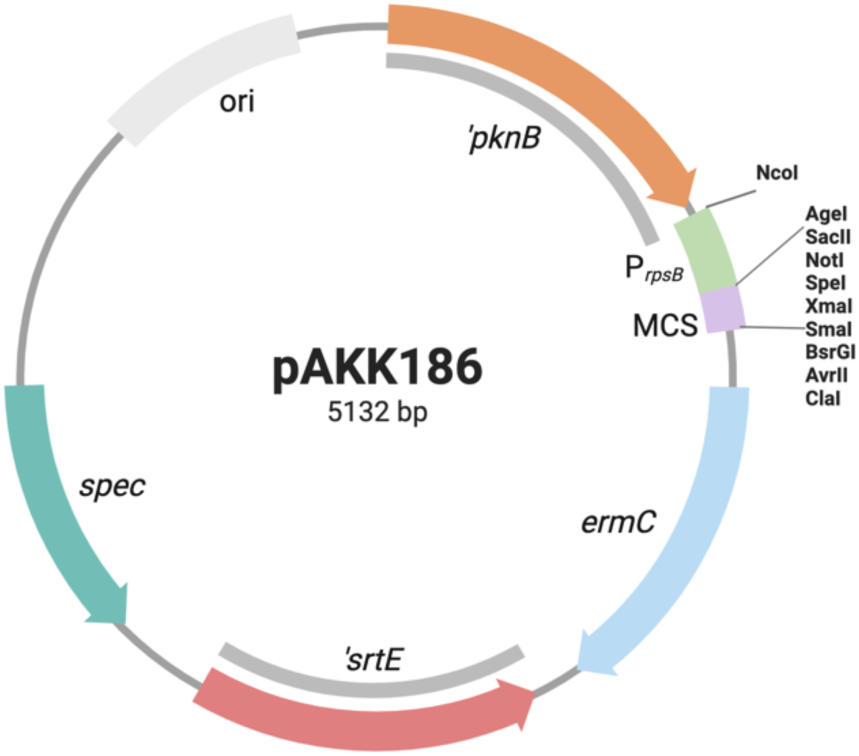
Complementation construct for *Gardnerella.* Regions of homology with the *G. vaginalis* ATCC 14018 chromosome are shown in gray. pAKK186 contains the *rpsB* promoter from ATCC 14018 for constitutive expression of a gene cloned into the multiple cloning site (MCS). Plasmid integrants will be Erm^R^ Spc^R^, while double recombinants will be Erm^R^ Spc^S^.

We PCR-amplified the wild-type *vly* gene, including its RBS, from ATCC 14018 and directionally cloned this sequence into pAKK186, resulting in pAKK187 (**Fig. 9A**). After treatment with HaeIII methyltransferase, we transformed the plasmid into the *vly* deletion mutant, AKK132. Five Erm^R^ transformants were patched to plates containing spectinomycin to determine whether the isolates had the plasmid integrated via a single crossover and would be Erm^R^ Spec^R^, or had undergone double-crossover recombination, which would result in Erm^R^ only. Two of the five isolates were resistant to both antibiotics, while three were resistant to Erm only. One of the Erm^R^ Spc^S^ isolates was analyzed by PCR to confirm that it had undergone double-crossover recombination and incorporated *vly* at the *pknB*-*srtE* locus (**Fig. 9B**). Spotting dilutions of the wild type, *vly* deletion mutant (AKK132), and complemented mutant (AKK136) on HBT agar showed that the mutant is non-hemolytic, while the complement exhibited the hemolytic phenotype like the wild-type strain (**Fig. 9C**). In quantitative assays, the wild type was strongly hemolytic and the *vly* mutant showed virtually no hemolytic activity on human red blood cells. The complemented mutant appeared only somewhat reduced in hemolysis activity compared to the wild type and was statistically not different from wild type (**Fig. 9D**).

**Figure 9.**
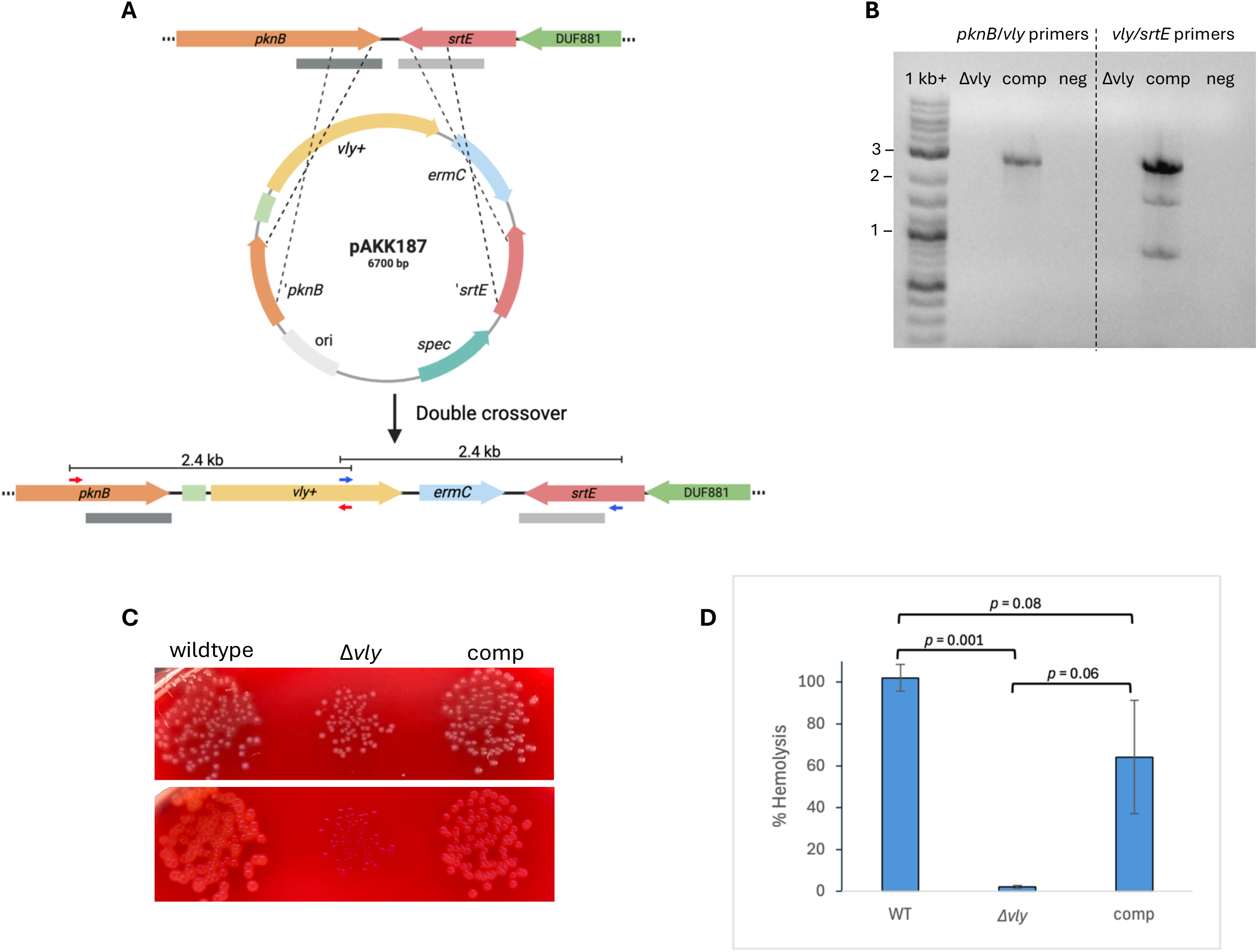
Complementation of *vly* mutant restores hemolysis activity. (A) Schematic depicting the complementation region in AKK132, construct pAKK187, and the double recombinant. pAKK187 contains *vly*+ and e*rmC* cloned between ‘*pknB* and ‘*srtE*. In the first step, transformants of AKK132 (Δ*vly*) containing the plasmid were selected for by plating on medium with erythromycin. Erm^R^ isolates were patched to plates containing spectinomycin to identify transformants that had either integrated the plasmid (Erm^R^ Spc^R^) or had undergone a double crossover (Erm^R^ Spc^S^), as depicted in the schematic. (B) PCR analysis confirming the presence of *vly* at the complementation site in a recombinant strain. Primer pairs (shown as red or blue arrows in (A) that anneal outside of the ‘*pknB* or ‘*srtE* fragment and within *vly* were used to amplify regions ∼2.4 kb in size. Lanes: 1kb+ = DNA ladder; Δ*vly* = mutant; comp = complemented mutant; neg = No DNA negative control. DNA sequencing confirmed the presence of *vly.* (C) Hemolysis phenotypes of wild type (ATCC 14018), Δ*vly* mutant (AKK132} and complemented mutant (AKK136) on HBT agar. The top panel shows the plate illuminated from ambient light. The bottom panel shows the plate illuminated by a light box from below. (D) Hemolysis assay of concentrated supernatants from cultures of small colony variants. The mean ± standard deviation of three independent assays is shown. Statistical significance was assessed by the Welch’s *t*-test for wild type versus Δ*vly* and Δ*vly* versus complement; and by the Student’s *t*-test for wild type versus complement.

### Mutagenesis in a different *Gardnerella* species, a sialidase deletion mutant in *G. pickettii*

To determine if our methods would work for other *Gardnerella* species, we deleted the sialidase gene *nanH3* in *G. pickettii* strain 3336 (*35*). We PCR-amplified two regions of the 3336 chromosome that were upstream or downstream of the *nanH3* gene and were each approximately 0.8kb. These DNA fragments were ligated into the positive/negative selection plasmid pAKK174 such that they created an in-frame deletion of *nanH3*, omitting 95% of the *nanH3* coding sequence. The resulting plasmid, pAKK191, was transformed into 3336, and Erm^R^ transformants were obtained at a frequency of 1x10^-6^. After overnight growth of one of these transformants without antibiotic, plating of the bacteria on BHI-YDS agar containing 4CP yielded isolates that had lost the plasmid insertion. PCR screening of seven isolates identified four that had lost the insertion and kept the *nanH3* deletion, and DNA sequencing verified the deletion. We used the chromogenic substrate X-Neu5Ac to assay for sialidase activity in the *Gardnerella* strains. Lysates of wild-type 3336 exhibited significant sialidase activity whereas little or no sialidase activity was present in lysates from the *nanH3* mutant (**Fig. 10A**). These results demonstrate that our mutagenesis methods and tools are not limited in their use to only *G. vaginalis* but also work for other *Gardnerella* species and generate mutations at a similar frequency.

**Figure 10.**
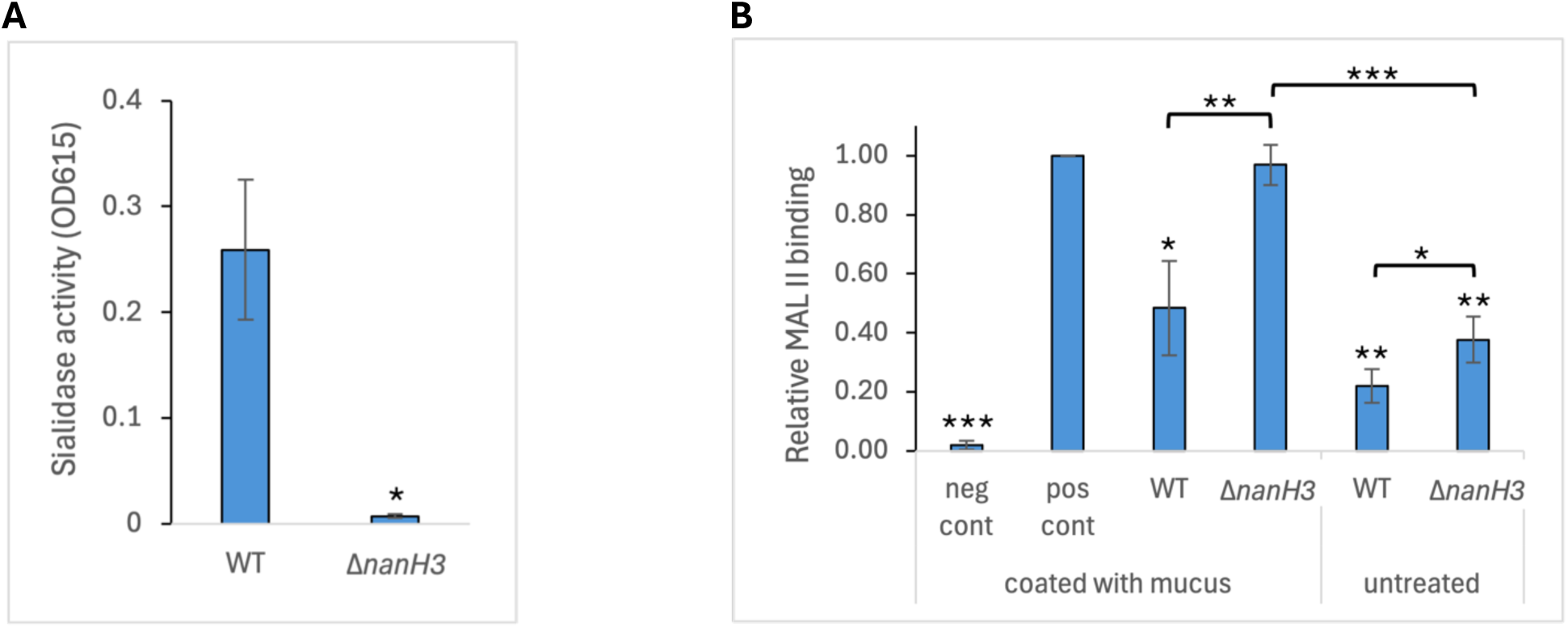
A *G. pickettii nanH3* mutant is deficient in sialidase activity and in cervical mucus degradation. (A) Bacterial lysates were mixed with the chromogenic substrate X-Neu5Ac and incubated overnight at 37°C. Absorbance at OD_615_ was measured. The mean ± standard deviation of three independent assays is shown. Statistical significance was assessed by the Welch’s *t*-test. (B) Wells of a microtiter plate were pre-coated with cervical mucus, followed by treatment with 3336 or the *nanH3* mutant. Lectin MAL II was used to quantify terminal sialic acids on mucus glycans. Controls included wells that did not receive MAL II (neg) and wells that received MAL II but no bacteria (pos). In addition, bacteria were applied to wells that had not been pre-coated with cervical mucus (untreated). Data were normalized to the positive control. The mean ± standard deviation of three independent assays is shown. Statistical significance was assessed by the Welch’s *t*-test for comparisons to the positive control, and by the Student’s *t*-test for comparisons between bacterial samples. Significance is shown relative to the positive control or between treatments as indicated by brackets. * *P* < 0.05, **, *P* < 0.01, ***, *P* < 0.001.

We examined the function of NanH3 by testing its ability to degrade human cervical mucus. Mucus was collected from human cervix tissue grown in organ culture. Cultures of wild type strain 3336 or the *nanH3* mutant were added to wells of a 96-well plate coated with cervical mucus and incubated for 24 hours to allow for mucus degradation. MALII lectin was added to bind remaining sialic acid residues in the cervical mucus, and the amount of MALII binding was measured. The wild type strain degraded more than 50% of the mucus while the *nanH3* mutant showed no significant reduction in sialic acid-containing mucus (**Fig. 10B**). Interestingly, controls containing 3336 or the *nanH3* mutant but without cervical mucus addition showed some lectin binding. This result indicates that either the bacteria have a sialic acid-containing molecule on their surface even in the absence of mucus or that the bacteria are able to bind the lectin directly. Overall, these results demonstrate that NanH3 acts in the degradation of human cervical mucus.

## Discussion

We have built a set of tools and methods for genetic manipulation of *Gardnerella* species. The methods allow for targeted mutagenesis and use selectable markers. We demonstrated that the constructs work for strains from two different *Gardnerella* species. Making plasmid insertions is straightforward, and maintaining antibiotic selection will allow for such insertion mutations to be stably maintained. Using the positive/negative selection plasmid to deliver point mutations or deletions to the chromosome was achieved. Such mutants are more desirable than insertions since they are stable and can be designed to have minimal polar effects on nearby genes. Furthermore, this method can be used iteratively so as to make multiple mutations in the same strain. The complementation construct was effective in delivering an expression construct to the chromosome. Hemolysis was restored in the *vly* mutant by expression of *vly* from this construct.

Getting the methods to work for transformation and produce antibiotic resistant *Gardnerella* isolates at a frequency substantially above the spontaneous antibiotic resistance frequency required overcoming several hurdles. Growth of *Gardnerella* in the presence of DCS significantly increased transformation frequency, likely due to affecting the barrier function of the envelope. DCS inhibits biosynthesis of both D-Ala and the D-Ala-D-Ala dipeptide used for making the UDP-MurNAc pentapeptide that is a precursor to peptidoglycan (*42*). The D-Ala-D-Ala portion of newly added peptidoglycan is used in the transpeptidation reaction for crosslinking peptidoglycan strands together (*47*). Thus, DCS-treated *Gardnerella* may have less cell wall or may have less-crosslinked cell wall. Another step that increased transformation frequency was the use of circular DNA. The reason for this difference is unclear, but it is possible that linear constructs were more subject to *Gardnerella* nucleases. Finally, a restriction barrier had to be overcome. Methylation of transforming DNA or knock-out of the *haeIIIR* gene encoding the HaeIII-like restriction enzyme in ATCC 14018 was effective in greatly increasing transformation. This last result is consistent with the results Kularatne and Hill observed when developing a method for transforming *G. vaginalis* with a *Bifidobacterium – E. coli* shuttle plasmid. Their shuttle plasmid was digested in vitro if treated with ATCC 14018 lysate but not with ATCC 49145 lysate, and they then used ATCC 49145 for their transformations (*48*). We found that plasmids with larger regions of homology to the *Gardnerella* chromosome yielded increased frequencies for insertion mutants, as was previously noted in *Neisseria gonorrhoeae* and *Streptococcus pneumoniae* (*49, 50*). Insert sizes of 500-1800 bp gave similar high transformation frequencies, but even a plasmid with an insert of 200 bp gave transformants. Overall, our methods improved targeted mutagenesis in *Gardnerella* from undetectable above background spontaneous resistance at 10^-9^ antibiotic resistant CFU/total CFU to approximately 10^-6^ antibiotic resistant CFU/total CFU for the recombinant strains.

We produced insertion mutations, a point mutation, and a deletion in the vaginolysin gene. Vaginolysin is a pore-forming toxin that lyses human red blood cells, epithelial cells, and neutrophils (*37, 51*). Vaginolysin binds human CD59, and this specificity likely explains the human-specific nature of *Gardnerella* infection (*37*). The role of vaginolysin in infection is debated. It has been proposed to act in nutrient acquisition, adherence, detachment from the cell surface, or destruction of neutrophils to facilitate immune evasion (*32, 52*). However, in a three-dimensional cell culture model, CD59 was found on the basolateral surface of cells, and vaginolysin added to the apical side failed to permeabilize the human cells (*53*). Others found that vaginolysin added to human vaginal or cervical cell lines resulted in membrane vesicle release from the human cells rather than lysis (*54*). In in vitro assays, all of our *vly* mutants lost hemolytic activity for human red blood cells. To test vaginolysin function with a more relevant tissue, we infected human cervix tissue in organ culture. Lactate dehydrogenase assays demonstrated that infection with wild-type *G. vaginalis* resulted in cell lysis in the cervix tissue, and infection with the vaginolysin mutant resulted in levels of cell lysis no different than uninfected control.

We produced a sialidase mutant of *G. pickettii* strain 3336. Sialidase activity is only found in some *Gardnerella* clades and species (*12, 55*). Sialidase is considered to be a likely virulence factor since sialic acid caps some glycan strands on mucins, preventing their degradation (*55*). Mutation of the *nanH3* gene in 3336 was sufficient to eliminate sialidase activity as measured with a chromogenic substrate. We further tested the function of NanH3 by measuring the ability of the mutant and wild-type strains to degrade human cervical mucus. The wild type was able to significantly reduce the amount of sialic acid-containing mucus present, while the *nanH3* mutant lost that ability. Sialidase is thought to degrade mucus for nutrient acquisition or to allow *Gardnerella* to access the host cell surface. *Gardnerella* can also use sialidase to desialylate immunoglobulin A, facilitating its degradation and preventing it from functioning in immune clearance of the bacteria (*32, 33*).

Limitations of our study include the limited number of species, strains, and variants that we attempted to transform and the limit in the types of mutations we attempted to create. We focused this study on the type strain of *G. vaginalis,* ATCC 14018, and on a sialidase producing strain of *G. pickettii,* strain 3336. We have not tried to transform strains from all of the 13 species of *Gardnerella*, nor have we measured transformation rates in additional strains in the species we did study. While we expect our methods to be generally applicable, there may be species or strains that are not transformable with our methods and constructs. A wide variety of restriction-modification systems are present among *Gardnerella* species and would limit transformation (*56*). We previously found that *Gardnerella* undergo colony phase variation (*35*). We have not attempted to determine if large colony variants are better or worse in transformation than small colony variants. Similarly, we did not compare opaque colony variants to translucent. We focused on making targeted insertions, point mutations, and deletions. Eventually, random mutagenesis will be needed to identify loci necessary for *Gardnerella* virulence, and methods for making transposon mutations or other random mutations throughout the chromosome will be needed.

New treatments are needed for BV. Antibiotic treatment only clears the infection in 60-70% of patients. Furthermore, many patients that are successfully cured, relapse within a number of months (*24*). To better treat BV, we need to identify the essential virulence factors. Using our mutagenesis methods and tools, *Gardnerella* mutants can be made and tested for loss of virulence in human cell or tissue infection in vitro or in the estradiol-treated mouse model (*35, 53, 57*). Identified virulence factors would serve as targets for antibiotics or be used in vaccines. Deletion mutants lacking genes for virulence factors, such as vaginolysin or sialidase, might be able to colonize the vagina without causing disease. Thus, vaginal microbiota transplants containing such nonpathogenic bacteria might be used after antibiotic treatment to occupy the niche and prevent reinfection with BV-causing bacteria. Furthermore, loss of factors that *Gardnerella* use to fight the immune system (such as sialidase and vaginolysin) might allow the adaptive immune system to develop protective responses that would block or clear any *Gardnerella* infection and thus prevent BV.

## Materials and Methods

### Bacterial strains, oligos, plasmids, primers, and growth conditions

Bacterial strains and plasmids used in this study are presented in Table S1. *Gardnerella vaginalis* ATCC 14018 was used to develop a method for electrotransformation starting with the methods for transformation with oligos described by Garcia (*40*). *Gardnerella* species exhibit phase variation in colony size (*35*); due to the faster growth rate of the large colony variant, we used the large morphotype for our studies. *G. vaginalis* was initially streaked from frozen stocks onto human blood tween (HBT) plates (Thermo Scientific^TM^) and grown overnight at 37°C with 5% CO_2_. To prepare electrocompetent cells, ATCC 14018 was grown in BHI-YDS media, which consisted of brain heart infusion (BHI) supplemented with 0.5% yeast extract, 0.2% glucose, and 0.1% starch. BHI-YDS agar plates were supplemented with 10% FBS and when necessary, antibiotics at the following concentrations: 50 or 100 μg/ml streptomycin (Str); 0.03 μg/ml erythromycin (Erm); 16 μg/ml spectinomycin (Spc); 8 μg/ml tetracycline (Tet). BHI agar supplemented with 10% FBS and 5% Fildes Enrichment, designated BHIFF medium, was used as nonselective medium for *G. vaginalis*. *E. coli* cultures were grown in LB media containing the appropriate antibiotic at the following concentrations: 500 μg/ml Erm; 100 μg/ml Spc. Primers, oligos, and gblocks (Table S2.) were purchased from Integrated DNA Technologies.

### Preparation of electrocompetent cells

*G. vaginalis* ATCC 14018 was streaked from a frozen stock onto an HBT agar plate and grown at 37°C with 5% CO_2_ overnight (24 hr). Colonies were swabbed from the HBT plate into 6-12 ml BHI-YDS medium, and the density was adjusted to an OD_600_ of 0.05. The culture was grown in a 15-ml conical tube held stationary overnight (18 hr). The overnight culture was diluted into 30-45 ml of fresh BHI-YDS medium to an OD_600_ of 0.1-0.15 and incubated statically at 37°C with 5% CO_2_. For initial transformation experiments, cells were harvested when the OD_600_ reached 0.35. In later transformations, cultures were grown to an OD_600_ of 0.25-0.3, then D-cycloserine (DCS) was added to a concentration of 250 µg/ml, and the culture was incubated another 1.5 hr before harvesting the cells. Cultures were placed on ice for 15 min and the cells were divided between two 50-ml Oak Ridge tubes and harvested by centrifugation for 10 min at 8000 rpm in a Sorvall RC6+ centrifuge at 4°C. The supernatant was removed, and each cell pellet was resuspended in 1 ml of ice-cold 10% glycerol in water and transferred to two 1.5-ml microfuge tubes. Using 1.5-ml microfuge tubes rather than Oakridge tubes at this step minimized cell loss. The cell suspension was centrifuged for 5 min at 8000 rpm at 4°C in a refrigerated centrifuge. The supernatant was discarded, and the cells were washed twice more. After the first centrifugation, the cell pellet was sticky and difficult to resuspend. However, with the successive washes, the pellet became easier to resuspend and formed a uniform suspension. The final cell pellets were combined and resuspended in a volume of ice-cold 10% glycerol that was 1/100 of the starting culture volume. Aliquots of 80 μl were placed in 1.5-ml microcentrifuge tubes on ice or stored at -80°C.

### Transformation protocol

Oligo DNA (10 μg), or plasmid DNA (1.5-2 μg), was mixed with 80 μl of electrocompetent cells and incubated on ice for 5 min. Cells without DNA were included as a negative control. The cell suspensions were transferred to pre-chilled 0.1-cm gap cuvettes (BioRad) and electroporation was performed, initially on a BioRad Micropulser unit with 2.2 kV, and later on a BioRad Gene Pulser or BioRad Gene Pulser Xcell unit with parameters of 200 Ω, 25 μF, and 2.2 kV. Time constants were recorded following the electrical pulse. Cells were immediately resuspended in 4 ml of pre-warmed BHI-YDS medium and allowed to recover statically overnight (18-20 hr) at 37°C with 5% CO_2_. Serial dilutions of the overnight cell suspensions were spotted on BHIFF agar to obtain total CFU/ml. Aliquots of the recovered cells were spread on BHI-YDS agar supplemented with the appropriate antibiotic. For transformations with oligos, 50 µl or 100 µl aliquots were plated; for transformations with plasmids, 1-ml aliquots that were concentrated 10-fold were plated. Plates were incubated at 37°C with 5% CO_2_ for two days. The transformation frequency was calculated as the antibiotic resistant CFU/ml divided by the total CFU/ml. Putative transformants were restreaked to BHI-YDS plates containing the appropriate antibiotic. Putative *rpsL* transformants were sequenced to verify incorporation of the mutation present on the oligo, while isolates transformed with plasmids were analyzed by PCR to verify the presence of the plasmid at the target locus in the chromosome.

### Methylation of plasmids

Plasmids were isolated from *E. coli* with the Qiagen Miniprep Kit, then methylated with HaeIII methyltransferase (NEB) according to the manufacturer’s instructions. Methylated plasmid was purified from the reaction mixture with the Qiaquick PCR Cleanup Kit (Qiagen) or the Monarch DNA and PCR Cleanup Kit (NEB), according to the manufacturer’s instructions. Approximately 1.5–2 μg of methylated plasmid was concentrated by centrifugation under vacuum in a speed vac to 1-2 μl in preparation for electroporation.

### Construction of insertion-duplication mutants

To construct an insertion-duplication (ID) plasmid for interrupting the gene for vaginolysin (*vly*), an approximately 0.5-kb internal fragment of *vly* was amplified by PCR from *G. vaginalis* strain ATCC 14018 with primers vly_SacI-F and Gv_abc-R. The PCR product was digested with SacI and ClaI and ligated into pIDN1 that had been digested with the same enzymes and dephosphorylated. The ligation mixture was transformed into chemically competent *E. coli,* and transformants were selected on LB plates containing Erm. The recombinant plasmid, pAKK134, was treated with HaeIII methyltransferase (NEB), and approximately 1.5-2 μg of methylated plasmid was electroporated into ATCC 14018 electrocompetent cells. Controls included cells without DNA, and cells with 10 μg of the 44-nt oligo rpsL4mut. Following overnight (18-20 hr) recovery, the negative control and transformed cultures were plated on selective agar. After 2 days of incubation at 37°C with 5% CO_2_, colonies were enumerated. Colonies from the plasmid transformation plates were restreaked to the same medium to confirm resistance to erythromycin. Erm^R^ isolates were screened by colony PCR with a primer that annealed to the genome (vly-F), along with a primer that annealed to the vector (pmob-R) to identify transformants that had the plasmid integrated at the correct locus. Three *vly* insertional mutants, AKK107-1/2/3, were characterized for hemolytic activity.

We constructed an insertion-duplication plasmid to interrupt the gene encoding the HaeIII-like endonuclease gene in ATCC 14018. We PCR-amplified a 447-bp internal fragment of the *haeIIIR* gene from ATCC 14018 with primers Gv_haeIII_SacI-F and Gv_haeIII_ClaI-R, and ligated the digested PCR product into the SacI/ClaI sites of pIDN1. This construct, pAKK139, was treated with HaeIII methyltransferase and transformed into ATCC 14018 as described for pAKK134. PCR analysis of Erm^R^ transformants with a primer that annealed to the genome (haeIII_screen-F) and one that annealed to the vector (pmob-R) confirmed plasmid integration at *haeIIIR*.

### VLY hemolysis assay

*Gardnerella* strains were streaked onto HBT plates from frozen stocks and grown overnight at 37°C with 5% CO_2_. Colonies was swabbed into BHI-YDS medium, and suspensions were adjusted to an OD_600_ of 0.05. Replicates of 200 µl of inoculum were placed in the wells of a 96-well tissue culture-treated plate and grown statically at 37°C with 5% CO_2_ for 21-24 hr. The initial and final ODs were measured as an indication of growth, and culture supernatant from triplicate wells was combined. Cells were removed by centrifugation, and 500 µl of supernatant was concentrated with a 30-kD cut-off column (Amicon Ultra 0.5 ml). Samples were stored at -80°C prior to analysis. Single donor human whole blood was purchased from Innovative Research Inc. Erythrocytes were collected by centrifugation (500x*g* for 5 min), plasma was removed, and the cells were washed three times with sterile PBS. A 1% solution of erythrocytes was prepared. Concentrated supernatant was serially diluted with PBS, and 100 µl of each dilution was mixed with 100 µl of the 1% erythrocyte suspension in a V-bottomed 96-well plate. Positive and negative controls for hemolysis were a 0.1% solution of Triton X-100 (Tx) and PBS, respectively. The plate was incubated for 30 min at 37°C with 5% CO_2_ and then centrifuged for 10 min at 2000 rpm to pellet intact erythrocytes. Supernatant was transferred to duplicate wells of a flat-bottomed 96-well plate and the absorbance (A) at OD_415_ was measured. Percent hemolysis was calculated as (A_sample_ – A_PBS_)/ (A_Tx_ – A_PBS_) X 100.

### LacZ insertion-duplication constructs

Four internal fragments of the beta-galactosidase gene (*lacZ*) from ATCC 14018 ranging in size from 0.2-1.8 kb were cloned into pDIN1. The 0.2-kb ‘*lacZ*’ fragment was ordered as a gblock from IDT (Integrated DNA Technologies). The 0.5-, 1- and 1.8-kb fragments were PCR-amplified from genomic DNA of ATCC 14018 with Gv_lacZ_SacI-F paired with Gv_lacZ(0.5kb)_XhoI-R, Gv_lacZ(1kb)_XhoI-R, or Gv_lacZ-R3, respectively. The gblock and PCR products were digested with SacI and XhoI, then cloned into the SacI/XhoI sites of pIDN1, resulting in constructs pAKK144, pAKK145, pAKK146, and pAKK148, which contain successively larger ‘*lacZ*’ inserts. The plasmids were treated with HaeIII methyltransferase and transformed into ATCC 14018 competent cells, resulting in insertion-duplication mutants AKK112, AKK113, AKK114, and AKK15, respectively. Integration at *lacZ* was confirmed by the white colony phenotype on BHI-YDS agar supplemented with 40 µg/ml of X-Gal.

### Plasmid excision assay

Strains AKK112-AKK115 were streaked out onto HBT agar and grown overnight at 37°C with 5% CO_2_. Colonies were swabbed from 24 hr plates into BHI-YDS medium and 6 ml of bacterial suspension was adjusted to an OD_600_ of 0.05. Liquid cultures were grown stationary at 37°C with 5% CO_2_ for 18-20 hr. Serial dilutions were prepared and spread on BHI-YDS agar containing 40 µg/ml X-Gal, while spot dilutions were plated on BHI-YDS agar to obtain total CFU/ml. After overnight incubation, blue colonies on the plates with X-Gal were enumerated and the frequency of plasmid excision was calculated as blue CFU/ml divided by total CFU/ml.

### Construction of positive-negative selection plasmids for creating unmarked mutations

To create unmarked mutations in *G. vaginalis*, we constructed an allelic exchange vector that contained a counter-selectable marker. In *E. coli*, PheS with the double mutation T251S A294G incorporates the toxic analog 4-chloro-DL-phenylalanine (4CP) into proteins, resulting in cell death (*58*). Alignment of the protein sequences of PheS from ATCC 14018 and *E. coli* K12 showed that ATCC 14018 contained the conserved Thr and Ala residues. Therefore, we designed a gblock containing the *pheS* from ATCC 14018 with the double mutation (T286S A333G), designated *pheS_mut2_*. We also incorporated a strong promoter (from the *gap* gene in ATCC 14018) to drive expression of *pheS_mut2_*. The gblock was digested with ClaI and XhoI and cloned into the ClaI/XhoI sites of pIDN1, creating pAKK163. Next a gblock with a 0.7-kb internal fragment of the vaginolysin gene that contained a nonsense mutation (E155*; GAA to TAA change) near the middle was digested with SacI and SmaI and ligated into the SacI/SmaI sites of pAKK163, resulting in plasmid pAKK165.

Another version of the positive-negative selection plasmid, pAKK174, was constructed that contained the *pheS_mut2_* gene with alternative codons (designated *pheS_mut2_alt_*) to reduce the possibility of integration at the *pheS* locus. The codon optimization tool by IDT was used to convert the *pheS_mut2_* codons from *G. vaginalis* to codons optimized for another member of the *Bifidobacteriaceae* family, *Bifidobacterium longum*. After the optimization, codons that appeared to be rare in the original ATCC 14018 *pheS* sequence were manually changed to more representative codons. Next a gblock was designed to create a complete deletion in *vly.* DNA fragments upstream (484 bp) and downstream (491 bp) of *vly* were fused to create a 1935-bp deletion that removes the entire *vly* coding sequence (1551 bp), in addition to some sequence flanking either side of the gene (277 bp and 107 bp, respectively). The Δ*vly* gblock was digested with SpeI and SmaI and cloned into the SpeI/SmaI sites of pAKK174, generating pAKK176.

### Creation of unmarked *vly* mutants using positive-negative selection constructs

A two-step method of gene replacement was used to create unmarked mutations in *vly* with pAKK165 (nonsense mutation) or pAKK176 (full deletion). First, ATCC 14018 was transformed with the methylated plasmid, and integration via a single crossover event was selected for by plating on medium with Erm. Integration at or near *vly* was confirmed by PCR analysis. Second, after growth of the integrant on medium without Erm, plasmid loss due to a second recombinational event was selected for by plating on BHI-YDS medium supplemented with 10% FBS and 1 mM 4CP. Isolates that have excised the plasmid do not incorporate the toxic 4CP into proteins and are therefore able to grow in its presence. If the second crossover occurs on the same side of the mutation that the first crossover occurred on, the wild-type gene is restored. However, if the second crossover occurs on the opposite side of the mutation, the mutation is incorporated into the genome. After 2 days, colonies were patched onto HBT for phenotypic screening. Sequencing confirmed the presence of the point mutation in or deletion of *vly*.

### Cervix infections

Human cervix samples were obtained from consented donors undergoing hysterectomies through the National Disease Research Interchange (NDRI). Donors were 50 years of age or younger and had not undergone chemotherapy or radiation treatment. Tissues were shipped overnight on ice in Dulbecco’s Modified Eagles Medium (DMEM). On the day of arrival, endocervical and ectocervical regions were processed into 3-mm biopsy punches. Tissue punches were allowed to recover overnight and were maintained in cervical tissue culture medium (CTCM: CMRL-1600 medium, 0.1 µg/ml hydrocortisone 21-hemisuccinate sodium, 1 µg/ml bovine insulin, 2 mM L-glutamine, 5% heat-inactivated FBS) with penicillin and streptomycin at 37°C with 5% CO_2_ until the infection experiment. This work was determined to be exempt as human subjects research by the University of Wisconsin Health Sciences IRB as NDRI codes each tissue sample and does not label or provide identifying information.

Endocervical and ectocervical punches were washed with sterile PBS and moved to fresh CTCM. ATCC 14018 and AKK124 were streaked onto HBT from freezer stocks and incubated overnight at 37°C with 5% CO_2_. The strains were restreaked onto HBT and incubated overnight. Bacteria were swabbed from the plates into PBS, washed in CTCM, and added at a final OD_600_ of 0.2 (∼1 x 10^8^ CFU/ml) in 1 ml of CTCM to cervix tissue in a 24-well plate. Two replicates per treatment were included. Plates were incubated at 37°C with 5% CO_2_ for 24 hr. 50 µl aliquots of supernatant were removed after 0, 12, and 24 hr of incubation, mixed with the Promega LDH Storage Buffer (200mM Tris-HCl (pH 7.3), 10% Glycerol, 1% BSA), and stored at -20 °C until they were analyzed for lactate dehydrogenase activity (Promega LDH-Glo™ Cytotoxicity Assay). The activity of LDH in the supernatant samples was measured using the bioluminescent assay according to the manufacturer’s protocol. The infections were repeated with at least three separate human donor tissues.

### Construction of complementation plasmid for *G. vaginalis*

We identified a region in ATCC 14018 that had convergent genes (*pknB* and *srtE*) that had high nucleotide conservation between clade 1 strains and thus could serve as a complementation site in *G. vaginalis*. We designed a gblock that had 0.8-0.9-kb fragments containing the 5’ ends of the *pknB* and *srtE*, with a multiple cloning site (MCS) between the two fragments. We also included the promoter from the 14018 *rpsB* gene to direct constitutive expression of the gene of interest. Inclusion of unique restriction sites on either side of the *rpsB* promoter (NcoI and AgeI) allows for replacement with a different promoter. The gblock had SacI and XhoI sites added to the ends to enable cloning into these same sites in pAKK182, an *E. coli* plasmid that contains a spectinomycin resistance gene from *Enterococcus faecalis*, that we found provided resistance in *G. vaginalis*. The *ermC* gene was subsequently cloned into the SalI site within the gblock, resulting in the complementation construct pAKK186.

We directionally cloned the *vly* gene from 14018 into the AgeI and BsrGI sites within the MCS, producing pAKK187. pAKK187 was methylated with HaeIII methyltransferase and electroporated into AKK132 (Δ*vly*), selecting for Erm^R^ transformants. Erm^R^ isolates were patched to plates containing spectinomycin to identify those that had integrated the full plasmid by single-crossover recombination rather than the desired transformants that had integrated only the region between *pknB* and *srtE* via double-crossover recombination. Some isolates were resistant to both Erm and Spec, while others were resistant to Erm only, suggesting these isolates had undergone a double crossover. PCR analysis was conducted to confirm that the Erm^R^ Spec^S^ isolates were double recombinants.

### Creation of *nanH3* deletion mutant in *Gardnerella pickettii*

A construct was made that contained *nanH3* from *G. pickettii* strain 3336 with a large in-frame deletion spanning amino acids 6-783 of the 812-amino acid protein. Two DNA fragments of approximately 0.8 kb were PCR-amplified from 3336. One fragment contained upstream flanking sequence and the first 5 codons of *nanH3*, followed by a SpeI site; the other fragment had a SpeI site followed by the last 30 codons of *nanH3* and downstream flanking sequence. The fragments were generated by PCR with primer pairs nanH3_up_SacI-F/nanH3_up_SpeI-R and nanH3_down_SpeI-F/nanH3_down_XhoI-R. The fragments were sequentially cloned into pAKK174 which resulted in construct pAKK191; the insert contained the Δ*nanH3* allele with a SpeI site inserted in-frame between the two fragments. Strain 3336 lacks the gene for the HaeIII-like endonuclease, so pAKK191 was not treated with the HaeIII methyltransferase prior to electroporation. Competent cells of 3336 were prepared with DCS treatment as described for ATCC 14018, and 2 µg of pAKK191 was electroporated with the same parameters that were used for ATCC 14018. Erm^R^ colonies were obtained at a frequency of 1.6x10^-6^. An Erm^R^ integrant, AKK140, was grown overnight without selection and plated on medium containing 1 mM 4CP to select for recombinants that had lost the plasmid insertion. A PCR screen of seven recombinants showed that four had the Δ*nanH3* allele. The mutant was designated AKK141. DNA sequencing confirmed the presence of the in-frame deletion.

### Sialidase activity of 3336 and *nanH3* deletion mutant

Strains 3336 and AKK141 (Δ*nanH3*) were grown overnight on HBT media. Colonies were swabbed into PBS, and the bacterial suspensions were adjusted to an OD_600_ of 0.5. Cells from 1.5 ml of each suspension were harvested by centrifugation and resuspended in 0.75 ml of 0.1 M sodium phosphate buffer, pH 5.8. The suspensions were sonicated on ice for 30 sec (1 sec on/1sec off) at 40% amplitude to lyse the cells. Cellular debris was removed by centrifugation, and 100 µl of supernatant were transferred to a new microfuge tube. 5 µl of 10 mM X-NeuNAc substrate (APExBIO) was added to the samples or to buffer and incubated overnight at 37°C with 5% CO_2_. The samples were transferred to a flat-bottomed 96-well plate, and the absorbance at OD_615_ was measured on a BioTek Synergy HT plate reader. The blank measurement was subtracted from the samples. Three independent assays were performed.

### Assessment of *Gardnerella* sialidase activity on cervical mucus

Mucus was harvested from cervical tissue obtained from NDRI and stored at -20°C. An aliquot of cervical mucus was mixed with an equal volume of 0.1% dithiothreitol (DTT) in PBS and placed on a shaker for 1 hr at room temperature. A Bradford assay was conducted to measure the protein concentration of the suspension. The suspension was adjusted to ∼150 µg/ml with PBS and 100 µl was applied to wells of a 96-well tissue culture plate. The plate was incubated at 4°C overnight to allow the mucus to adhere to the wells. 3336 and AKK141 (Δ*nanH3*) were grown overnight on HBT, then swabbed into NYCIII media lacking glucose and adjusted to an OD_600_ of 0.2. Excess mucus was removed, and the wells were washed once with PBST (PBS + 0.05% Tween 20). 100 µl of the bacterial suspensions or medium alone were added to the wells and the plate was incubated at 37°C with 5% CO_2_ for 24 hr to allow for mucus degradation. The following day bacteria or medium were aspirated, and the wells were washed three times with 200 µl of PBST. To quantify remaining sialic acid, 100 µl of a 0.5 µg/ml solution of biotinylated *Maackia amurensis* Lectin II (MAL II) (Vector Laboratories) in PBS were applied to the wells and incubated at room temperature with gentle rocking for 1 hr. MAL II binds alpha-2,3 linked sialic acid. The wells were washed three times with PBST. 100 µl of HRP-conjugated streptavidin (R&D Systems), diluted 1:200 in PBS, were applied the wells and incubated at room temperature for 1 hr with rocking. Wells were washed three times with 200 µl of PBST, then 100 µl of TMB substrate (3,3’,5,5’ tetramethylbenzidine) were added to the wells and incubated 5 min at room temperature. 50 µl of 2N sulfuric acid were added to stop the reaction and the color was quantified by measuring the OD_450_ on a BioTek Synergy HT plate reader. Controls included wells that were pre-coated with cervical mucus but not treated with bacteria; the positive control had MAL II applied, while the negative control had PBS applied instead. Additional controls included bacterial suspensions that were added to wells that had not been pre-coated with cervical mucus.

### Statistical analysis

The F-test was used to compare variances between two groups. Statistical analysis was performed using a Student’s *t* test for data with equal variance or a Welch’s *t* test for data with unequal variance. *P* values of less than 0.05 were considered statistically significant.

## ACKNOWLEDGEMENTS

This work was supported by NIH grants R21AI182550 (JPD), R21AI153534 (JPD), and T32AI055397 (EMG). JPD and KKJ conceived of the study and interpreted results. AKK and EMG designed and performed the experiments, and interpreted results. AKK and JPD wrote the paper, and all authors approved of the final version. Materials in this study are the subject of a pending patent, and these materials are available through a material transfer agreement.

