## Supplementary material for "Genetic transformation of *Gardnerella* species and characterization of vaginolysin and sialidase mutants": Table S1

**Table S1. Bacterial strains and plasmids used in this study.**

| Strain or plasmid | Characteristics | Source/reference |
| --- | --- | --- |
| Bacterial strains |  |  |
| *Gardnerella* spp. |  |  |
| ATCC 14018 | *G. vaginalis* LacZ^+^, hemolytic, sialidase^-^ | ATCC |
| 3336 | *G. pickettii* LacZ^-^, hemolytic, sialidase^+^, Tet^R^ | M. Roberts |
| AKK107 | 14018 x pAKK134 (‘*vly*’) ID* mutant, Erm^R^ | This work |
| AKK110 | 14018 x pAKK139 (‘*haeIII*’) ID mutant, Erm^R^ | This work |
| AKK112 | 14018 x pAKK144 (0.2 kb ‘*lacZ*’) ID mutant, Erm^R^ | This work |
| AKK113 | 14018 x pAKK145 (0.5 kb ‘*lacZ*’) ID mutant, Erm^R^ | This work |
| AKK114 | 14018 x pAKK146 (1 kb ‘*lacZ*’) ID mutant, Erm^R^ | This work |
| AKK115 | 14018 x pAKK148 (1.8 kb ‘*lacZ*’) ID mutant, Erm^R^ | This work |
| AKK123 | 14018 x pAKK165 (*vlymut* *pheS*_mut2_) integrant, Erm^R^ 4CP-sensitive | This work |
| AKK124 | 4CP-resistant Erm^S^ AKK123, *vly* point mutant | This work |
| AKK131 | 14018 x pAKK176 (∆*vly* *pheS*_mut2_alt_) integrant | This work |
| AKK132 | 4CP-resistant AKK131, ∆*vly* mutant | This work |
| AKK136 | AKK132 x pAKK187 (∆*vly*/*vly*+) complemented mutant | This work |
| AKK140 | 3336 x pAKK191 (∆*nanH3* *pheS*_mut2_alt_) integrant, Erm^R^ | This work |
| AKK141 | 4CP-resistant AKK141, ∆*nanH3* mutant | This work |
| *E. coli* |  |  |
| TAM1 | Used for cloning | RapidTrans^TM^ |
| TOP10 | Used for cloning | Invitrogen^TM^ |
| Plasmids |  |  |
| pIDN1 | *E. coli* cloning vector, Erm^R^ | (*49*) |
| pSPUC | *E. coli* cloning vector, source of spectinomycin resistance gene, Spc^R^ | (*59*) |
| pAKK134 | 0.5 kb ‘*vly*’ in pIDN1, Erm^R^ | This work |
| pAKK139 | 0.45 kb ’haeIII’ in pIDN1, Erm^R^ | This work |
| pAKK144 | 0.2-kb ‘*lacZ*’ in pIDN1, Erm^R^ | This work |
| pAKK145 | 0.5-kb‘*lacZ*’ in pIDN1, Erm^R^ | This work |
| pAKK146 | 1-kb *lacZ*’ in pIDN1, Erm^R^ | This work |
| pAKK148 | 1.8-kb ‘*lacZ*’ in pIDN1, Erm^R^ | This work |
| pAKK163 | counter selectable construct 1; *pheS*_mut2_ gblock in pIDN1, Erm^R^ | This work |
| pAKK165 | ’*vlymut*’ (point mutation) gblock cloned into pAKK163, Erm^R^ | This work |
| pAKK174 | counter selectable construct 2; pheSmut2_alt gblock cloned into pIDN1, Erm^R^ | This work |
| pAKK176 | ∆*vly* gblock cloned into pAKK174, Erm^R^ | This work |
| pAKK182 | Spec cassette cloned into XhoI/NsiI sites of pIDN1 (replacing *ermC*), Spc^R^ | This work |
| pAKK185 | ‘*pknB*_‘*srtE* gblock cloned into pAKK182, Spc^R^ | This work |
| pAKK186 | complementation plasmid for *G. vaginalis*; *ermC* cloned into pAKK185, Spc^R^ Erm^R^ | This work |
| pAKK187 | *vly*+ cloned into pAKK186, Spc^R^ Erm^R^ | This work |
| pAKK191 | ∆*nanH3* from 3336 cloned into pAKK174, Erm^R^ | This work |

*ID = insertion-duplication mutant

**Table S2. Sequences of primers, oligos and gblocks used in this study*.**

| Primers | Sequence 5’-3’ |
| --- | --- |
| vly_SacI-F | CTCTGAGCTCCTCAAATTGTGAACTTCAAGC |
| Gv_abc-R | TGAACGCTCAACACTTGC |
| vly-F | CATATGAAGAGTACAAAGTTCTACC |
| pmob-R | AACAGCTATGACCATGATTACGCCAAG |
| vly_screen-F | CGTCAAGGATAACGAAGTAGC |
| vly_screen-R | GCTACTTCGTTATCCTTGACG |
| Gv_haeIII_SacI-F | GATAGAGCTCGGATCTAACTGGCAAGAGA |
| Gv_haeIII_ClaI-R | GCTCATCGATCAAAGGTCTGTCACCATC |
| haeIII_screen-F | AGGCATTGGACTGCAAACTGG |
| Gv_lacZ_SacI-F | CTATGAGCTCGCATCATGGCTAGAGGATC |
| Gv_lacZ(0.5kb)_XhoI-R | TGATCTCGAGCACATCGGAAACCATGTCA |
| Gv_lacZ(1kb)_XhoI-R | GTATCTCGAGTCTGGGTTGTTTGACACG |
| Gv_lacZ-R3 | TGTAGCGTATTGTCGTCG |
| spec_ClaI-F | AGATATCGATTAACGTGACTGGCAAGAG |
| spec_NsiI-R | GACAATGCATACAGCTATGACCATGATTACG |
| vly_SpeI-F | CTATACTAGTCTGTAACTTCCGCAGTGA |
| vly_mid-F | TTCGATGCAGTACACAAGG |
| vly_up_screen-F | TGTGTGATGCGTGACCTTC |
| pknB_screen2-F | GCTGGAGAAGTTGTAGTTCC |
| srtE_screen3-R | GACCAGAACAGGTTGAGG |
| nanH3_up_SacI-F | CATAGAGCTCAGCACGCAGTAAATAGACC |
| nanH3_up_SpeI-R | GCATACTAGTCGCTGTTCCAATCATTATATTCC |
| 3336nanH3_down_SpeI-F | CAGTACTAGTGGAATCAACACAACTATTCCTTATGC |
| 3336nanH3_down_XhoI-R | TTAACTCGAGAACTAAGATGCCCTCGGT |
| Oligos |  |
| rpsL2mut | CGTGTGTACACCACCACTCCT**CG**GAAGCCAAACTCTGCGCTTCG |
| rpsL4mut | CGTGTGTACACCACCACTCC**ACGC**AAGCCAAACTCTGCGCTTCG |
| rpsL2mut_long | CGTGGCGTGTGCACTCGTGTGTACACCACCACTCCT**CG**GAAGCCAAACTCTGCGCTTCGTAAGGTTGCTCGTGTGCGCCTCAGC |
| rpsL4mut_long | CGTGGCGTGTGCACTCGTGTGTACACCACCACTCC**ACGC**AAGCCAAACTCTGCGCTTCGTAAGGTTGCTCGTGTGCGC CTCAGC |
| gblocks |  |
| 0.2 kb ‘lacZ’ gblock | CTATGAGCTCGCATCATGGCTAGAGGATCAAGATTATTGGAGACTACACGGCATTTTCCGTTCCGTTGAACTGTGTGCACATCCTTCCACCCATGTATCAAATCTTCATGTTGATGCAGATTATAGCAATGATACAAATACTGGGAAACTCGCATTCAGAGCAAACATTGAAGGAGACAATTTAAAAGATATTACACTACACGCTTACATCTCGAGGTAT |
| pheSmut2 gblock | CTGAATCGATCGTAATGCGTCGTACATACCGCACATTAAAATGAACATGTTAGTTTAAATAACGAAGACCAAAACGTATTGTTAAAAATCGCGAAAAATCGCTAGAACTATGCGTTAAGGTTTCACTCAAGGGAGATCCCATATGCTTCACAAAAGTACACATAACACGGATAGGAAGGGTGCTGTGGCAAGTAGTAAGCCGTTCGACGCGAAGGCAATTACAAAGGTAGTAAAAGAGGGTATAGCTTGTGTTCAAGCCGCTAAAACAATGGAAGAGTTGAAAGCGGCAAAAACTAAGTATGCAGGAGCTCAGTCTGCAATGACCCTTGCCAGCAAGTCTATTGGAAGTCTTCAAGTAGAAGAAAAGAAAGAAGCTGGCAAGATAATGTCGGCTTTACGCGCTGATTTTGGTCGTGAATTTGCAGCAGCCCAAGAGCGTATTAAGGCTATTGAAGAAGCTAATATGCTTCAAAAAGAAACAGTCGATATGACCTTGCCAATAAATCGTAAGCCGCTTGGCGCTCGCCATCCGATTGCGCGTATTATTGAAGACTTTGAAGACTTCTTTGTGTCTATGGGTTGGCAGATTTCTGCAGGACCAGAGGTAGAAACAGAGTGGTTCGATTTTGACGCGTTGAATTTTGGTCCAGATCATCCTGCTCGTCAAATGCAAGATACTTTTTATGTGCAAGGTAATCAGGCAAAGGATGCTGCAGGATTTGTTGGATCTAACATGGTTTTGCGTACGCAAACTTCTTCCGATCAGGTTCGTGCGTTAATTGAGCGAGGAGTACCTCTGTATATTGCGTCCCCAGGCCGAGTATTCCGCACGGATGAGCTTGATGCAACGCATACTCCAGTCTTCCACCAGTGCGAAGCTTTGGCAGTAGATAAACATTTGAGCATGGCTGATTTAAAGGGTGTTCTTGATCGCCTTGCTGTTGCAATGTTTGGTCCAGATGCAAAAAGCCGTTTGCGCCCAAGCTACTTCCCGTTC**T**C**T**GAGCCTAGTGCGGAGTTGGATCTTTGGTTCCCAGATAAGAAGGGTGGCCCAGGTTGGATTGAATGGGGCGGATGCGGAATGGTGAATCCGAATGTTCTTAAGTCTGCAGGCTTAGACCCAGATGTTTATACGGGATTTG**GA**TTCGGTGTTGGCTTGGAGCGCACTTTGCTGCTTCGTCACGATATTAACGATATGCACGATTTGGTTGAAGGCGATAAGCGCTTCAGTGAACAGTTTGTGATGGGAGAGTAGCTCGAGTATG |
| vlymut gblock | CTGTACTAGTGAACCAGCTACATCTTGCGCAGCTAAGAAAGACTCGTTGAATAATTATTTGTGGGATTTGCAATACGATAAAACAAACATTCTCGCCCGTCATGGCGAAACCATTGAGAACAAATTCTCCAGCGACAGCTTCAACAAGAACGGTGAATTCGTTGTTGTTGAGCATCAGAAGAAGAACATCACCAATACAACTTCAAATTTGTCGGTTACTTCCGCCAACGATGATCGCGTATACCCAGGTGCTCTTTTCCGTGCTGATAAGAATTTGATGGACAATATGCCAAGCCTGATTTCTGCAAACCGCGCTCCAATAACGTTGAGCGTTGATTTGCCGGGATTCCACGGCGG**T**AAAGTGCTGTAACTGTTCAGCGCCCAACCAAGAGTTCTGTAACTTCCGCAGTGAACGGCTTAGTTTCTAAGTGGAATGCACAATATGGAGCAAGTCATCATGTTGCAGCTCGCATGCAGTACGATTCTGCAAGCGCACAAAGCATGAACCAGCTCAAGGCTAAGTTTGGTGCTGATTTTGCCAAGATTGGTGTTCCGCTGAAGATTGATTTCGATGCAGTACACAAGGGTGAGAAGCAGACTCAAATTGTGAACTTCAAGCAAACTTACTACACCGTAAGCGTTGATGCACCAGATAGCCCAGCAGATTTCTTTGCTCCTTGCACTACGCCAGACAGCTTGAAGAACCGTATCGATTGAG |
| pheSmut2_alt gblock | GTTACTCGAGCGTAATGCGTCGTACATACCGCACATTAAAATGAACATGTTAGTTTAAATAACGAAGACCAAAACGTATTGTTAAAAATCGCGAAAAATCGCTAGAACTATGCGTTAAGGTTTCACTCAAGGGAGATCCCATATGTTGCATAAATCTACGCACAACACTGACCGCAAAGGAGCCGTGGCGTCTAGCAAGCCGTTCGACGCTAAAGCTATTACTAAAGTTGTGAAAGAAGGTATAGCCTGCGTACAGGCAGCTAAAACAATGGAAGAGTTGAAGGCGGCTAAGACGAAGTATGCCGGCGCTCAATCCGCTATGACACTTGCAAGCAAAAGCATTGGATCCTTGCAAGTTGAGGAGAAGAAGGAAGCTGGTAAGATTATGTCTGCTTTGCGCGCGGATTTTGGCCGCGAGTTCGCTGCAGCCCAAGAGCGCATTAAAGCGATTGAAGAGGCTAATATGCTTCAAAAGGAAACGGTAGATATGACATTGCCGATAAACCGCAAACCGCTTGGAGCACGTCATCCGATTGCGCGTATTATTGAAGATTTCGAGGACTTCTTCGTGTCTATGGGATGGCAAATTTCTGCGGGTCCGGAAGTTGAGACGGAGTGGTTTGATTTCGACGCGTTGAATTTCGGTCCAGACCACCCAGCTCGTCAGATGCAGGATACCTTCTACGTACAAGGAAATCAGGCCAAAGATGCGGCTGGATTTGTTGGATCGAATATGGTGCTTCGTACACAGACGTCTTCTGATCAAGTTCGTGCATTGATTGAACGTGGTGTTCCACTTTATATTGCGAGCCCGGGCAGGGTGTTCAGGACTGACGAACTTGACGCGACTCACACCCCAGTGTTTCATCAATGCGAGGCGCTTGCTGTGGATAAGCATCTTTCCATGGCCGACCTTAAAGGCGTTCTTGATAGGCTTGCTGTTGCAATGTTCGGCCCAGACGCGAAGTCTCGTCTGCGCCCGTCTTACTTTCCGTTTA**GC**GAACCGTCTGCGGAATTGGACCTTTGGTTTCCGGACAAAAAAGGAGGACCAGGCTGGATAGAATGGGGTGGATGCGGCATGGTGAACCCAAATGTACTTAAATCCGCTGGTCTTGACCCAGACGTTTACACGGGATTTG**GT**TTTGGTGTCGGTCTTGAACGCACACTGTTGCTGAGGCATGACATTAACGATATGCATGACTTGGTAGAGGGCGACAAACGTTTCTCTGAGCAGTTTGTTATGGGTGAGTAGGGTACCTCAG |
| ∆*vly* gblock | GTCGACTAGTCCAGACTCACTTCTAGGAGCGCCAGAAAGCGCTTCTTCTTGAGTCAATGGCGTAGCAGGCTCGTCTGGGCGCAAATACGCTTCAAGAGGTACTGGATTTGCAGTAGGTGGCACATCTTCGCCAGCAACTTCCTGCACTTTGTTAATGGATTCTGCAATGACGTTTAATTCGCCTTGCAAGCGACTGATTTCTTCATCGCTTAAAGCAATCTGGGACAAAACACCCAAATGCTCAATTTCTTCGCGTGTGAATGTAGGCATAACCTCAACTATATGTGTGATGCGTGACCTTCTTATAGAAGAAAAAAGCAATATTGATAAAAGATTTACACACGTTACTGTAGAAATATTTTCCCAATTTTAGAACATGATTGTCGATCCATATTTTCTAGGGAAAATTATGTGTAATATTAAAAAAAATTTTATTAGTGAGACAATAAGCGAAATAACAATTATTGTCACATACAGTTTTCATAAAATTAACGTTATTTGCAGAATATTTAAATATTTTAGATATCGCGCTAAATATCGCGATGCTTTTCCACAACGTGGCCAATTGCATACATGACTACGCCCCAAGCAAGAACCACCAAGCCTGATTGCCACCATGTAAAAACATATGCGTTTGGTGGAAGCTGTGCACCCGAATTAGGGGAGCCGCCCAAGAATTTCTCCACCGCTGTGGCTGGCAAAAGCTGTATAAGTATTGAATTCCACTTCGCGAAATTGCTTGCAAACATAATAATGCTAAGAACACTAGGCAAAATCATCACGGCTCCAATAACGCACATAATTCCGCCAGCAGTTGACTTGCAAATCATGCCAAAGCCGTACGCCATTGCTGCTACAACAACCATAATTGCAGGAGAGCCAAGAAACAGCGTAAGAGGCAATTTCCACGCGTTGCTTCCAGATAATCCAGAAGTGTTGTCTCCTATAAAGGCTAATTCTGCAGCGCCTAGCGAAACTGCCATTGCCCGGGTATA |
| Gv pknB_srtE gblock | ATATGAGCTCGCTCTTGGTCTTATCCCAGATATTTTGGAAGATGATAAATCTTCTCAACCAGAAGGAACTTTTACAAAACAGCTTCCTAAAGGTGGAGCTAAAGTTTCTTCAGGATCTAGAGTAAGCGTATGGTTCTCTGTAGGTCCACAGTCTACTAAGATTCCAGATGTTACTGGTAAATCGCAAGATGTAGCACGTAAAGCTCTTGAACGTGCTGGATTTAAGATATCTAATGTTCGTGTAGAAGATAGCACGGAAGTTAAGAAGAATCATGTTACTCGTACAGATCCTTCTGCAGATTCGTTTGCAGACAAGGGGTCTATGGTTACCTTATACATTTCTTCTGGTCTTACAAAGATCCCTGATGGTTTAGTTGGACAATCTAAGGATGTTGTAACAAGCGAATTGCAAAATCTTGGTTTTACTGTAAATGTTGTTGAGGAAGAATCTGATACTGCATCTGAAGGAACTGTGACCAAGATGAACCCATCTTCTGGTGCTGCAGTAAAGCCTCATAGCTCTGTAACAGTATATGTTTCTAAAGGAAAGCCTAAAGTTGAAGTTCCTCGTTTGGCTGTTGGCACTGTAACTTTCAAGCAGGCTAAACAAGTATTGGAAGCAAAAGGCTTTAAGGTTGTTGCATCTGATGCATCAGCTAAAGATGATGACATAGTTACTGGAATGTCTGAAAAAGAAGGAGCAAAGATTGACAAAGGTTCAACAATCACTTTGACTGTGAAATCTGCTACTCCTCCATCGGACCTTACAAAGCCAGGTACGGGATTAGATTTGAATAAACCTTCTACTGAAGCCCCATCTACATCTACTACAACAACAGATACAGGTTTGTAAAAAGCAAAGCATAAAGCCAGTAAGTTTAATAAGCTTACTGGCTTTCCATGGTTAGTATTAAGCGTCTGATATGATTATCTTTTGCGTGTCCATTGAGCATGAATGTTGTCGCGTTATATGATTGCTCGTTTTTTATTGTCTACTCGGATACGCGAATACGCGCCATGTACGCGTCCGTTCGTGTCGGTCGGCTTGGATTTAGACATATGTTATCCAAGTTGCATGCGAACAGGCATGGCCAGGCAAGAACCGACACCGGTCTCCACCGCGGTGGCGGCCGCACTAGTGGATCCCCCGGGCTGCAGGAATTCTGTACACCTAGGATATCAAGCTTATCGATACCGTCGACGGCCGGCCTTTATTATTTATTAATCTTATCTACTTACTTATTATCTACTTACCCCTTAGTAAGGCGTTGCTGTGTATGCAGACATTTCTCTTAAGAATGGTATATTTGCCGAAACAAACGGGTATACCCATTGCATAAGTATCATAATGCATATTAGGTAGATAATTAAGGAGAGAATTAATCTCACTGGAAGAATTCCAGGCTGTAGCCTCATGAGTCCGCCAAAGATGCTGAATTCAGGCTTTGGCTTTAGACCTGCTTTAATCTCGCGTCTTAATGGCCACTGCCATGCAATAGCTCCAGCAGCAAAGATAATCAAATAAACTATCAGCGCGCCAATCATAAGCGGAACAAGCGAATCGAGATGAGAGATCAAAGATTGTTGCTCGTTATTAACAAACTTAACTTTTCCGTTTTCATCTAAAGTAGAAAGCTCTTTTGGAATACCGTCAGAAACTTTAGCCCAATAGTCAAGCTCTCCAAAGCTAACAAAGCGGAACTTAGGTGTAGAGAACTTAGGTTCGCATGTAATAATGGTGATCATGCGTTTCTTTGGCTCTTTACTATGGTTTATTGGATCTGGGTCAAGAACCTCAACCTGTTCTGGTCTAACAATCTTGTGAGTAATGTATTTGTAAACGAACCAATAATCTTTAGTTCGAAGAATAATAGAATCTCCCTTTTGGAACCTATCAACATCCGCAAGAGGCTGACCATAACCATTGCGGTGACCAATAATTGTAATGTTTCCGATTTCTCCAGGCAACTCAGTTTTAGGATAATGACCTAATCCTCGACGATTAAGTATCTCCATATCTACACCTTCAATAACATTACGCTCCCATTGATCACCGAAGCGTGGAATATAGATTCGTGCTCGAGTATC |

*Restriction sites are underlined. Bolded nucleotides in oligos and gblocks represent changes to the wild-type sequence.
